# Parental kynurenine 3-monooxygenase genotype in mice directs sex-specific behavioral outcomes in offspring

**DOI:** 10.1101/2024.12.18.629210

**Authors:** Snezana Milosavljevic, Maria V. Piroli, Emma J. Sandago, Gerardo G. Piroli, Holland H. Smith, Sarah Beggiato, Norma Frizzell, Ana Pocivavsek

**Author notes:** Corresponding Author: Ana Pocivavsek, Ph.D., University of South Carolina School of Medicine, Department of Pharmacology, Physiology and Neuroscience, Building 1, D26, 6311 Garners Ferry Rd, Columbia, SC 29209, USA.

## Abstract

**Background:** Disruptions in brain development can impact behavioral traits and increase the risk of neurodevelopmental conditions such as autism spectrum disorder, attention-deficit/hyperactivity disorder (ADHD), schizophrenia, and bipolar disorder, often in sex-specific ways. Dysregulation of the kynurenine pathway (KP) of tryptophan metabolism has been implicated in cognitive and neurodevelopmental disorders. Increased brain kynurenic acid (KYNA), a neuroactive metabolite implicated in cognition and sleep homeostasis, and variations in the *Kmo* gene, which encodes kynurenine 3-monooxygenase (KMO), have been identified in these patients. We hypothesize that parental *Kmo* genetics influence KP biochemistry, sleep behavior and brain energy demands, contributing to impairments in cognition and sleep in offspring through the influence of parental genotype and genetic nurture mechanisms.

**Methods:** A mouse model of partial *Kmo* deficiency, *Kmo* heterozygous (HET-*Kmo^+/−^*), was used to examine brain KMO activity, KYNA levels, and sleep behavior in HET-*Kmo^+/−^* compared to wild-type control (WT-Control) mice. Brain mitochondrial respiration was assessed, and KP metabolites and corticosterone levels were measured in breast milk. Behavioral assessments were conducted on wild-type offspring from two parental groups: i) WT-Control from WT-Control parents, ii) wild-type *Kmo* (WT-*Kmo^+/+^*) from *Kmo* heterozygous parents (HET-*Kmo^+/−^*). All mice were C57Bl/6J background strain. Adult female and male offspring underwent behavioral testing for learning, memory, anxiety-like behavior and sleep-wake patterns.

**Results:** HET-*Kmo^+/−^* mice exhibited reduced brain KMO activity, increased KYNA levels, and disrupted sleep architecture and electroencephalogram (EEG) power spectra. Mitochondrial respiration (Complex I and complex II activity) and electron transport chain protein levels were impaired in the hippocampus of HET-*Kmo^+/−^* females. Breast milk from HET-*Kmo^+/−^* mothers increased kynurenine exposure during lactation but corticosterone levels were unchanged. Compared to WT-Control offspring, WT-*Kmo^+/+^* females showed impaired spatial learning, heightened anxiety, reduced sleep and abnormal EEG spectral power. WT-*Kmo^+/+^* males had deficits in reversal learning but no sleep disturbances or anxiety-like behaviors.

**Conclusions:** These findings suggest that *Kmo* deficiency impacts KP biochemistry, sleep behavior, and brain mitochondrial function. Even though WT-*Kmo^+/+^* inherit identical genetic material as WT-Control, their development might be shaped by the parent’s physiology, behavior, or metabolic state influenced by their *Kmo* genotype, leading to phenotypic sex-specific differences in offspring.

**Plain English summary:** Interactions between genetic and environmental factors are carefully regulated during the intricate process of brain development. While genetic information is directly inherited from parents, emerging evidence suggests that parental genetic factors can also shape the environment influencing children’s development in a sex-specific ways. Disruptions in brain development can impact cognitive and behavioral traits and increase the risk of neurodevelopmental conditions such as autism spectrum disorder, attention-deficit/hyperactivity disorder, schizophrenia, and bipolar disorder. This study explored how *kynurenine 3-monooxygenase* (*Kmo*) genotype affects female and male mice, focusing on potential sex-specific behavioral changes in offspring born to parents with a genetic disruption in *Kmo*. We found that female and male mice with partial *Kmo* deficiency experienced reduced sleep and increased sleep pressure. In female mice, *Kmo* deficiency impaired mitochondrial energy production in the brain. We also observed alterations in tryptophan metabolism and nutrient composition in the breast milk of *Kmo*-deficient females. In adult offspring born to *Kmo*-deficient parents, females exhibited learning difficulties, heightened anxiety-like behaviors, and sleep disturbances. In contrast, male offspring showed mild cognitive impairments but no major sleep issues. These findings highlight that parental *Kmo* genotype can influence sex differences in cognitive and sleep-related behaviors in offspring. This underscores the importance of considering parental genetic factors when studying neurodevelopmental disorders and associated behavioral outcomes.

**Highlights:**

- Mice with a partial *Kmo* deficiency, *Kmo* heterozygous (HET-*Kmo^+/−^*), were used to evaluate the impact of parental *Kmo* genetics on kynurenine pathway biochemistry, brain mitochondrial function, sleep patterns, and cognitive performance.
- Female and male HET-*Kmo^+/−^* mice exhibited elevated brain KYNA levels and had reduced sleep, prolonged wakefulness, and altered spectra power during sleep. In HET-*Kmo^+/−^* females, hippocampal mitochondrial respiration and electron transport chain protein levels were significantly altered. Breast milk from HET-*Kmo^+/−^* females increased kynurenine exposure during lactation to offspring, including their wild-type (WT-*Kmo^+/+^*) offspring.
- Sex differences were observed in learning and memory, anxiety-like behavior, and sleep-wake patterns between WT-Control, from wild-type parents, and WT-*Kmo^+/+^* offspring from HET-*Kmo^+/−^* parents. Female WT-*Kmo^+/+^* offspring had impaired spatial learning, increased anxiety-like behavior, reduced sleep, but elevated delta power during sleep. Male WT-*Kmo^+/+^* offspring exhibited deficits in reversal learning only and not sleep impairments.

## BACKGROUND

Neurodevelopmental disorders, including autism, attention-deficit/hyperactivity disorder (ADHD), schizophrenia, and bipolar disorder have a strong genetic component and are highly heritable. Genome-wide complex analyses estimate over 70% heritability of neurodevelopmental disorders [1–4]. Single nucleotide polymorphism (SNP) studies suggest shared genetic factors contribute to the overlap in heritability across these disorders [5]. Recent genome-wide association studies (GWAS) highlight increasing evidence of genes influencing multiple traits, emphasizing the need for research to unravel the functional impacts of genetic variations [6, 7]. At the same time, clinical studies suggest that neurodevelopmental processes are strongly shaped by sex-influenced biological mechanisms, which significantly affect behavioral outcomes [8, 9]. For example, autism and ADHD are more common in males [10], while females experience more severe schizoaffective symptoms [11]. However, the role of sex-related mechanisms in parental genetic contribution to the offspring neurobiological outcomes remain poorly understood.

Neuroactive metabolites of the kynurenine pathway (KP), the primary route of tryptophan metabolism (**Figure 1A**), are disrupted in patients with autism, ADHD, schizophrenia and bipolar disorder [12–20] and a role for the KP in the etiology of neurodevelopmental illnesses has been highlighted in various rodent models [21–24]. Kynurenine 3-monooxygenase (KMO) converts kynurenine to 3-hydroxykynurenine (3-HK) (**Figure 1A**), and ultimately regulates production of the ubiquitous co-factor nicotinamide adenine dinucleotide (NAD^+^)[25]. Human studies point to a causal relationship between intronic *Kmo* SNP, rs2275163, and neurocognitive deficits frequently reported in patients with schizophrenia and psychotic bipolar disorder [26, 27], and postmortem brain tissue analysis suggest an association between *Kmo* SNPs and reduced KMO activity [27, 28]. When KMO activity is reduced, excessive amounts of kynurenine are converted to kynurenic acid (KYNA) by kynurenine aminotransferases (KATs) (**Figure 1A**). Intronic *Kmo* SNP, rs10158645, and exonic *Kmo* SNP, rs1053230, have been associated with increased cerebrospinal fluid levels of KYNA in patients with schizophrenia and bipolar disorder [29, 30]. Elevated levels of the brain KYNA, an endogenous inhibitor of alpha 7 nicotinic acetylcholine (α7nACh) and N-methyl-D-aspartate (NMDA) receptors, contribute to impairments in sleep and cognitive behaviors including spatial learning and cognitive flexibility [31–37]. These characteristics align with endophenotypes exhibited by individuals with cognitive and neurodevelopmental disorders [38–40].

**Figure 1.**
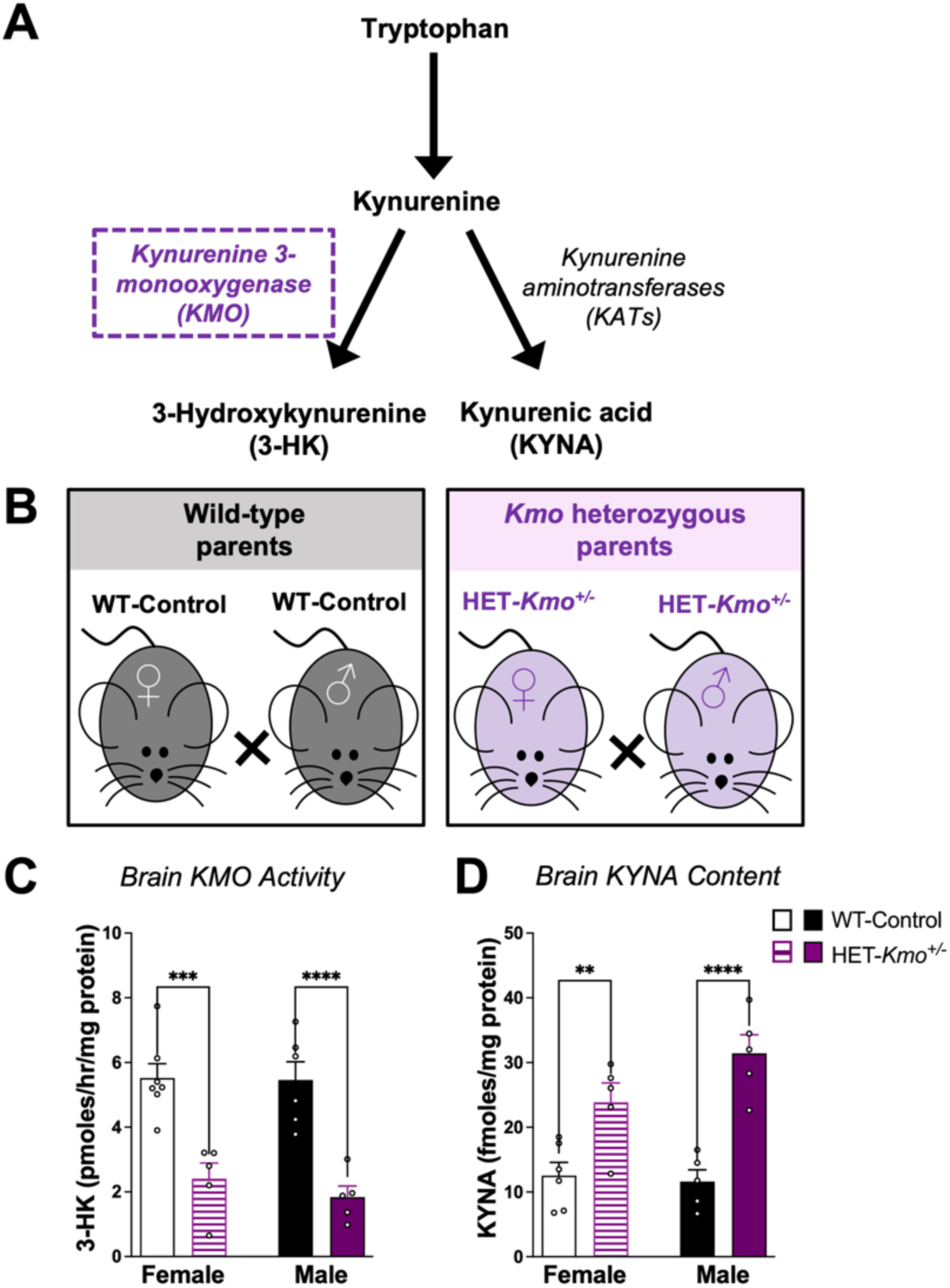
Decreased KMO activity and increased KYNA in the brain of *Kmo* heterozygous female and male mice. **(A)** Schematic representation of the kynurenine pathway denoting the pivotal enzyme KMO (in purple color) that is genetically modified in our present mouse model. **(B)** Schematic representation of the experimental parental groups. Wild-type control (WT-Control) breeding pairs and heterozygous *Kmo* (HET-*Kmo^+/−^*) breeding pairs. **(C)** Brain KMO activity in HET-*Kmo^+/−^* mice is reduced compared to WT-Control mice (Unpaired t test ***P<0.001, ****P<0.0001). **(D)** Brain KYNA levels in HET-*Kmo^+/−^* mice are increased compared to WT-Control mice (Unpaired t test **P<0.01, ****P<0.0001). Data are mean ± SEM. N = 5-7 per group.

Perinatal factors, including environmental exposure to stress and dietary challenges – included elevated maternal exposure to kynurenine – are linked to impaired neurodevelopment and long-term behavioral outcomes in offspring [24, 41, 42]. In rodent models, maternal dietary elevations in kynurenine result in increased KYNA in the fetal compartment [21, 42, 43], leading to impairments in neurochemistry, sleep, and cognitive behavioral outcomes in offspring [24, 44–46]. Specifically, in utero, a disproportionate increase in KYNA is observed in the fetal brains from dams heterozygous for the *Kmo* gene (HET-*Kmo^+/−^)* compared to control mice [21].

Based on this, we hypothesize that offspring from HET-*Kmo^+/−^* mice exhibit distinct and enduring behavioral endophenotypes. Reduced brain KMO activity and elevated KYNA levels were observed in HET-*Kmo^+/−^* mice, along with disruption in sleep parameters, including duration, architecture, and spectral power. Sleep-wake disturbances may elevate brain energy demands, tightly regulated by mitochondrial respiration [47–49]. Since the KMO protein is localized in the outer mitochondria membrane [50] and may influence mitochondrial function [51], we examined mitochondrial respiration in HET-*Kmo^+/−^* mice and observed alterations in hippocampal mitochondrial function. To explore maternal contributions, we analyzed for the first time KP metabolite levels in the breast milk of female HET-*Kmo^+/−^* mice, which may influence offspring brain development and behavior [52]. Based on these findings, we hypothesized that the unique biochemical profile of HET-*Kmo^+/−^* mothers contributes to behavioral outcomes in their offspring and analyzed behavioral phenotypes and sleep in two cohorts: wild-type C57Bl/6J (WT-Control) mice and wild-type offspring of HET-*Kmo^+/−^* parents (WT-*Kmo^+/+^*). While both groups shared the same genotype, female WT-*Kmo^+/+^* offspring exhibited significant deficits in sleep and cognitive behavior, underlying sex-specific impairments in *Kmo* genetic nurture. Finally, we performed behavioral assessments on adult offspring, including spatial learning and memory, anxiety-like behavior, and sleep-wake behavior. WT-*Kmo^+/+^* offspring from HET-*Kmo^+/−^* parents exhibited sex-specific deficits compared to controls, suggesting that parental *Kmo* deficiency disrupts neurodevelopment and imposes adverse behavioral outcomes in offspring.

## METHODS

### Animals

Wild-type (WT-Control) C57Bl/6J mice were purchased from Jackson Laboratory (Bar Harbor, ME, USA) and bred in-house, along with a transgenic mouse line featuring a targeted deletion of the *Kmo* gene on the C57Bl/6J background [53]. Nomenclature for all mice used in the study is defined as the following:

1. **WT-Control:** Wild-type C57Bl/6J mice used as breeding pairs, in parental studies, and as adult offspring in behavioral studies.
2. **HET-*Kmo^+/−^*:** Mice heterozygous for *Kmo*, used as breeding pairs and in parental studies.
3. **WT-*Kmo^+/+^*:** Wild-type offspring from HET-*Kmo^+/−^* breeding pairs, used as adult offspring in behavioral studies.

Parental pair nomenclature is detailed in **Figure 1B**, and offspring nomenclature is shown in **Figure 5A**.

Adult mice used in experiments (2-4 months) were maintained in a temperature- and humidity-controlled facility fully accredited by the Association for Assessment and Accreditation of Laboratory Animal Care (AAALAC). Mice had free access to food and water on a 12-hour light-dark cycle. Lights were on at Zeitgeber time (ZT) 0 and lights off at ZT 12. All experimental protocols were approved by the Institutional Animal Care and Use Committee (IACUC).

### Genotyping

Offspring from HET-*Kmo^+/−^* parents were genotyped for presence or absence of the Pgk:neomycin resistance (NeoR) cassette [53]. DNA was extracted from a tail snip obtained at weaning on postnatal day (PD) 21 using standard method for nucleic acid purification by phenol chloroform extraction [54]. The concentration of DNA was measured via spectrophotometry (BioPhotometer Plus, Eppendorf, Enfield, CT, USA). DNA was amplified using polymerase chain reaction (PCR). For each DNA sample, one PCR verified the presence of the NeoR cassette and targeted disruption of the *Kmo* gene, and another PCR determined the absence of the NeoR cassette and the existence of the entire *Kmo* genomic locus. Primer sequences included: NeoR forward primer (CCTCGTGCTTTACGGTATCGCCGCTC), WT forward primer (TTCACCAGGTTGGGTAAAGC), and reverse primer (ATGCCTGCAACAACAATCAA) (Thermo Fisher Scientific, Waltham, MA, USA). The initial DNA denaturing in the PCR reaction was for 3 minutes at 94°C, followed by 35 cycles consisting of 30 seconds denaturing at 94°C, 30 seconds annealing at 67-69°C (a temperature gradient was used to determine the optimal temperature for each set of primers), and 80 seconds extension at 72°C. DNA fragments were separated by agarose gel electrophoresis (MyGel Mini Electrophoresis System, Accuris Instruments, Edison, NJ, USA) based on their size and visualized using ultraviolet light (UVP High-Performance UV Transilluminators, Thermo Fisher Scientific, Waltham, MA, USA).

### Breast Milk Collection

Breast milk was collected from N=10 WT-Control and N=11 HET-*Kmo^+/−^* mothers on PD 10-11. To allow for the mothers’ mammary glands to engorge with milk, pups were physically separated from mothers in a plastic box located in their parents’ cage for 5-6 hours, while still allowing visual, auditory, and olfactory interaction between mother and pups through transparent walls and perforated lid. Mothers and pups were reunited for at least 5 minutes until maternal behaviors were exhibited and nursing initiated. The mother was removed from the pups and lightly anesthetized with isoflurane (induction 3%; maintenance 0.5-1%). The experimenter pressed gently on the base of the teat moving upwards beginning on the inguinal mammary glands to stimulate milk expression [55]. Milk drops at the tip of the teat were quickly collected via a glass capillary tube and expelled into a 0.2 mL tube kept on ice. Milk expression was continued on abdominal, thoracic, and cervical mammary glands. Milk samples were frozen and stored at −80°C until biochemical analyses.

### Biochemical Analysis

#### Brain KMO activity and 3-HK measurement

KMO enzyme activity was evaluated in the brain tissue of N=13 WT-Control and N=10 HET-*Kmo^+/−^* adult mice, as previously described [21, 56]. Briefly, brain tissue samples were diluted 1:25 (w/v) and homogenized in 100 mM Tris-HCl buffer (pH 8.1) containing 10 mM KCl and 1 mM EDTA. 80 μl of diluted sample was incubated in a solution containing 1 mM NADPH, 3 mM glucose-6-phosphate, 1 U/mL glucose-6 phosphate dehydrogenase, 100 μM L-kynurenine, 10 mM KCl and 1 mM EDTA in a total volume of 200 μl for 40 min at 37°C. 50 μl of 6% perchloric acid was added to stop the reaction. To obtain blanks, 100 μl of the KMO inhibitor Ro 61-8048 was added to the incubation solution. Samples were centrifuged (16,000 x g, 15 min) and 20 μl of the supernatant were applied to a 3 μm C18 reverse phase high pressure liquid chromatograph (HPLC) column (HR-80; 80 mm × 4.6 mm; Thermo Fisher Scientific, Waltham, MA, USA) using a mobile phase containing 1.5% acetonitrile, 0.9% trimethylamine, 0.59% phosphoric acid, 0.27 mM EDTA, and 8.9 mM sodium heptane sulfonic acid at a flow rate of 0.5 mL/min. 3-HK (retention time ~11 minutes) was detected electrochemically (HTEC 500 detector)(Eicom, San Diego, CA, USA) in the eluate.

#### Brain and breast milk tryptophan, kynurenine, and KYNA measurement

Tryptophan, kynurenine, and KYNA were measured in biological samples by HPLC [21, 45]. Briefly, samples were diluted (brain 1:5 w/v; breast milk 1:50 v/v) in ultrapure water. Eighty μl of diluted sample was acidified with 20 μl of 25% perchloric acid and centrifuged (12,000 x g, 15 min). Twenty μl of the supernatant was isocratically eluted through a ReproSil-Pur C18 column (4mm x 100 mm; Dr. Maisch GmbH, Ammerbuch, Germany) using a mobile phase containing 50 mM sodium acetate and 3-5% acetonitrile (pH adjusted to 6.2 with glacial acetic acid) at a flow rate of 0.5 mL/min. Zinc acetate (500 mM) was delivered after the column at a flow rate of 0.1 mL/min. Tryptophan (excitation: 285nm, emission: 365nm, retention time ~13 minutes), kynurenine (excitation: 365nm, emission: 480nm, retention time ~7 minutes), and KYNA (excitation: 344nm, emission: 398nm, retention time ~12 minutes) were detected fluorometrically (ACQUITY UPLC H-Class PLUS System)(Waters Corporation, Bedford, MA, USA) in the eluate. Data were integrated and analyzed using Empower 3 software (Waters Corporation).

#### Corticosterone measurement in the breast milk

Breast milk samples were diluted 1:40 (v/v) with steroid displacement reagent and corticosterone levels were assessed using an enzyme-linked immunosorbent assay (ELISA) kit according to the manufacturer’s instructions (Enzo Life Sciences, Farmingdale, NY, USA). ELISA plates were read with a Synergy 2 Gen 5 microplate reader at 405 nm absorbance (BioTek Instruments Inc., Winooski, VT, USA).

### Sleep Studies

#### Surgical Procedure

Surgical implantation of the telemetry transmitter (HD-X02; Data Sciences International (DSI), St. Paul, MN, USA) with electroencephalographic (EEG) and electromyographic (EMG) electrodes in adult mice was conducted as previously described [57]. Briefly, following anesthesia with isoflurane (induction 5%; maintenance 1-2%), animals were placed in a stereotaxic frame (Stoelting Co., Wood Dale, IL, USA) to secure the head. Carprofen (5 mg/kg, s.c.) was administered at the start of the surgical procedure as an analgesic. A transverse incision was made on the dorsal abdominal surface for intraperitoneal implantation of the telemetry transmitter. Another longitudinal incision was made along the midline of the head and neck to exposure the skull and neck muscle. Two EMG leads were inserted directly into the dorsal neck muscle approximately 1.0 mm apart and secured into place. Two surgical stainless-steel screws (P1 Technologies, Roanoke, VA, USA) were implanted into two drilled burr holes (0.5 mm diameter) at 1.9 mm anterior/ +1.0 mm lateral and 3.4 mm posterior/ −1.8 mm lateral relative to bregma. Each EEG lead was wrapped around a screw and anchored with dental cement (Stoelting Co.). The dorsal incision was sutured, and the skin along the cement cap was reinforced with Gluture (Zoetis Inc, Parsippany, NJ, USA). Animals were singly housed and recovered postoperatively for at least 14 days prior to the start of EEG/EMG data acquisition.

#### Sleep Data Acquisition and Analysis

Sleep data were acquired at a continuous sampling rate of 500 Hz with Ponemah 6.10 software (DSI) in a designated room (48 hour recording period in male mice; 96 hour recording period in female mice to span the estrous cycle). Total number of subjects included N=14 WT-Control; N=14 HET-*Kmo^+/−^*; N=16 WT-*Kmo^+/+^*.

Digitized signal data were scored offline with NeuroScore 3.0 (DSI). EEG/EMG waveforms were hand-scored by visual inspection of 10-seconds epochs into one of three vigilance states: wake (low-amplitude, high-frequency EEG combined with high-amplitude EMG), NREM (high-amplitude, low-frequency EEG combined with low-amplitude EMG), or REM (low-amplitude, high-frequency EEG combined with very low EMG tone). Data were analyzed in 4-hour or 12-hour time bins for each vigilance state, assessing total duration, number of bouts, and average bout duration. NREM and REM sleep onset, and the number of transitions between vigilance states were also evaluated. The DSI module designed for periodogram powerbands in NeuroScore was used for NREM and REM sleep power spectra analysis in each phase. The Discrete Fourier transform (DFT) estimated the EEG power spectrum for defined frequency bandwidths: delta (0.5-4 Hz), theta (4-8 Hz), alpha (8-12 Hz), sigma (12-16 Hz), and beta (16-20 Hz). Relative home cage activity was reported by NeuroScore and evaluated in 12-hour time bins.

### Mitochondrial Respiration

#### Respiration parameters in isolated mitochondria

To measure respiratory parameters in mitochondria isolated from brains of N=5 WT-Control and N=5 HET-*Kmo^+/−^* female mice, we used a Seahorse XFe24 Analyzer following a previously published protocol with minor modifications [58]. For mitochondria isolation, mice were euthanized by decapitation following deep isoflurane anesthesia; brains were removed, and the cortex and hippocampus were dissected. Tissues were homogenized in 9 volumes of cold MSHE buffer (75 mM sucrose, 225 mM mannitol, 1 mM EGTA, 5 mM HEPES, pH 7.2) added 0.2% fatty acid free bovine serum albumin (BSA), using a Teflon glass homogenizer. Homogenates were centrifuged in 1.5 mL Eppendorf tubes at 800 x g at 4°C for 10 min, and the supernatants were passed to fresh tubes and centrifuged at 8,000 x g at 4°C for 10 min. The supernatants from the second centrifugation were discarded and the pellets containing mitochondria were resuspended in MSHE without BSA and centrifuged again at 8,000 x g at 4°C for 10 min. The final pellets were resuspended in a minimal volume of MSHE without BSA, and the protein content of the suspension was determined using the bicinchoninic acid (BCA) assay prior to diluting them in 1X MAS buffer (70 mM sucrose, 220 mM mannitol, 10 mM KH_2_PO_4_, 5 mM MgCl_2_, 1 mM EGTA, 2 mM HEPES, pH 7.2) with added respiratory substrates (10 mM succinate + 2 μM rotenone (to assess Complex II-dependent respiration), or 10 mM glutamate + 3 mM malate (to assess Complex I-dependent respiration).

Twenty μg of mitochondrial protein in 50 μl were loaded in two Seahorse XFe 24-well plates (one for succinate + rotenone dilutions, the other one for glutamate + malate dilutions), and the plates were centrifuged at 2,000 x g at 4°C for 20 min. Then, 400 μl of warmed (37°C) 1X MAS buffer containing the corresponding respiratory substrates were carefully added to the wells and the plates were incubated in a non-CO_2_ incubator at 37°C for 30 min. During the centrifugation of the plates, two Seahorse cartridges were loaded with the compounds to be injected during the experiment [Port A: 100 mM adenosine diphosphate (ADP), Port B: 40 μM oligomycin, Port C: 50 μM carbonyl cyanide 4-(trifluoromethoxy)phenylhydrazone (FCCP), Port D: 30 μM Antimycin A; all compounds get diluted 1:10 after the injection], and the cartridges were incubated in a non-CO_2_ incubator at 37°C for 30 min. The experiment started loading the cartridge for calibration, followed by the plate containing the mitochondria. The oxygen consumption rate (OCR) was then measured in basal conditions, and after the sequential port injections described above. Then, we subtracted the residual respiration obtained after the injection of antimycin A from the OCR traces to calculate the basal respiration (before any injection), state 3 (after ADP injection), state 4o (after oligomycin injection), and state 3u (after FCCP injection). The results were expressed in pmol O_2_/min/μg protein.

#### Western Blotting

Mitochondrial fractions prepared for Seahorse experiments were analyzed for the expression of representative complex I-V components by immunoblotting. Twenty μg of mitochondrial protein were separated in 12% polyacrylamide gels and transferred to nitrocellulose membranes. Blots were blocked with 5% non-fat dry milk in 50 mM Tris-HCl, 150 mM NaCl, pH 7.6 (TBS), then incubated with a total OXPHOS rodent western blot antibody cocktail mix (Abcam AB110413, 1:15,000) prepared in 2% non-fat dry milk in TBS containing 0.2% Tween 20 (TBST) overnight at 4°C with shaking, followed by four 5 min washes with TBST and a 60 min incubation with a secondary goat anti-mouse antibody labeled with a fluorescent tag (IRDye 680RD, LiCor 926-68070, 1:30,000) at room temperature. A second set of four 5 min washes with TBST was performed, followed by a final wash with TBS prior to image capture with a LiCor Odyssey CLX System running under iStudio. Results were normalized to the signal of VDAC2 (GTX104745, Gene Tex Inc., 1:5,000), using a donkey anti-rabbit antibody labeled with a fluorescent tag (DyLight SA5-10044, Invitrogen, 1:10,000). We also detected in the same samples the degree of 4-hydroxy-2-nonenal (HNE) modification of proteins using an anti-HNE antibody (HNE-11S, Alpha Diagnostics, 1:4,000) and the same donkey anti-rabbit secondary antibody listed above (1:8,000). Color images obtained with the LiCor system were converted to black and white and the intensity of the bands was quantified with ImageJ.

#### Protein Assays

Proteins in the samples analyzed by HPLC were evaluated using previously described Lowry method [59]. Total protein quantification for the mitochondrial respiration and Western Blotting experiments was performed by the BCA assay according to the manufacturer’s instructions (Thermo Fisher Scientific, Waltham, MA, USA).

### Behavioral Assays

Adult female and male mice used in the behavioral testing were 2-4 months old. Mice were brought into the behavior testing room at least 30 minutes prior to the start of the first trial of each experimental day to acclimate to the room. All behavioral tests were performed by an experimenter blind to the genotype condition.

#### Barnes Maze Paradigm

The Barnes maze paradigm was utilized to assess spatial learning and memory [60] in N=40 WT-Control and N=40 WT-*Kmo^+/+^* adult mice. Barnes maze consisted of an elevated circular platform (92cm diameter; 94cm elevation), with 20 holes around the edge (5cm hole diameter). One hole led to an escape box while the other 19 holes had shallow false bottoms. Extra-maze spatial clues were located on each wall of the designated behavior testing room. One day prior to the start of learning trials, each mouse was habituated to the maze by placing a mouse inside of the escape box for 120 seconds to become accustomed to the escape box, followed by allowing maze exploration for up to 180 seconds, and entrance to the escape box. Once inside the escape box the mouse was given 15 seconds inside then returned to its home cage. Spatial learning trials were performed for 3 consecutive days with 2 trials per day and a 4-hour inter-trial interval. Mice were placed in the center of the maze and allowed 180 seconds to locate the escape box. Once inside the escape box each mouse was given 15 seconds then returned to its home cage. On the 4^th^ day, the reversal trial was performed by rotating the Barnes maze 180 degrees. Barnes maze trials were acquired with EthoVision XT 15 video tracking system (Noldus, Leesburg, VA, USA). Distance traveled, errors committed, time spent immobile on the maze, and search strategy were assessed [45].

#### Elevated Zero Maze

The elevated zero maze (EZM) behavioral paradigm was utilized to assess anxiety-like behavior in N=44 WT-Control and N=30 WT-*Kmo^+/+^* adult mice. EZM is an elevated circular platform (50cm diameter; 62cm elevation), with two open/bright quadrants and two closed/dark quadrants arranged in alternate order. Mice, placed at alternating boundaries between open and closed areas, explored the maze for 300 seconds and overhead video tracking (EthoVision XT 15) was used for data acquisition. Number of entries to open and closed areas and the duration spent in open and closed areas were assessed [61].

#### Splash Test

Splash test was utilized to assess motivational behavior and self-care in N=44 WT-Control and N=25 WT-*Kmo^+/+^* adult mice. Individual mice were placed in fresh cages and then splashed on the dorsal coat with 400 µL of a 10% w/v sucrose solution at distance of 5 cm. Overhead video camera acquired behavioral data for 5 minutes following the splash. Manual scoring of grooming (cleaning of the coat) was obtained by two independent scorers using EthoVision XT 15 software. Grooming frequency and latency to groom were analyzed [62, 63].

#### Parental Care Behavior

Parental care behavior in N=7 WT-Control and N=10 HET-*Kmo^+/−^* breeding pairs was evaluated multiple times between PD 1 and PD 15 with up to 2 trials per day performed between ZT 2 and ZT 5 (early light phase) or ZT 6 and ZT 9 (late light phase). The experimenter entered the designated room at least 10 minutes before the observations period began and mice were not handled during the observations but rather remained undisturbed in their home cages. For 30 minutes the experimenter manually recorded the presence of the following categories of behavior every 3 minutes for a total of 10 observations per cage per trial: positive parental behaviors (pups in nest, nursing, grooming pups), self-care behaviors (eating and drinking, self-grooming), and negative parental behaviors (parent out of nest, pups out of nest, nest maintenance)[64, 65]. Of note, across observation periods, we never observed pups out of the nest.

## STATISTICAL ANALYSIS

Normality of data was assessed with Shapiro-Wilk test and data were visually inspected using Q-Q plots to confirm a relative bell-shaped distribution and the absence of outliers. All statistical analyses were performed using GraphPad Prism 9.0 software (GraphPad Software, La Jolla, CA, USA). Our current study design was statistically powered to allow for evaluation of sex as a biological variable. Significant interaction between sex and *Kmo* genotype was determined for several analyzed behavior parameters (**Supplementary Statistical Data**).

### Biochemical experiments

The brain tissue 3-HK and KYNA levels, as well as breast milk tryptophan, kynurenine, and corticosterone measurements were assessed by unpaired t test.

### Sleep studies

Sleep-wake architecture data were analyzed independently for each light and dark phase. Comparison between WT-Control and WT-*Kmo^+/+^* genotype groups was performed by a two-way repeated measures (RM) analysis of variance (ANOVA). Data inspection in 4-hour time bins for sleep-wake architecture parameters was evaluated with genotype as between-subject factor and time as within-subject factor. When sleep-wake data were compared between WT-Control and HET-*Kmo^+/−^* genotype groups, individual parameters were assessed separately for each vigilance state (REM, NREM, wake) by unpaired t test. Two-way RM ANOVA was performed for analysis of NREM and REM sleep power spectra where genotype (WT-Control and WT-*Kmo^+/+^*, or WT-Control and HET-*Kmo^+/−^*) was taken into consideration as between-subject factor and frequency as within-subject factor. Impact of genotype on relative cage activity, transitions between vigilance states, and sleep onset was determined by unpaired t test.

### Mitochondrial respiration

For respiration and Western Blotting experiments data were analyzed by unpaired t tests between the two genotypes.

### Behavioral experiments

Two-way RM ANOVA with genotype (WT-Control, WT-*Kmo^+/+^*) as between-subject factor and learning day as within-subject factor was used to evaluate Barnes maze parameters. Effect of genotype on Barnes maze search strategies was evaluated by contingency chi-square test for each day separately. Barnes maze reversal trial parameters, EZM parameters, and splash test parameters were evaluated by unpaired t test. Parental care behaviors were evaluated by two-way ANOVA with genotype as the between-subject factor.

Where appropriate, post hoc multiple comparison analysis between experimental groups is denoted within figure legend. Statistical significance was defined as P < 0.05.

## RESULTS

### Kynurenine pathway disruptions in the brain of *Kmo* heterozygous mice

Compared to WT-Control mice, KMO activity was decreased in the brain of both female and male HET-*Kmo^+/−^* mice compared to WT-Control mice, confirming a loss of function in brain KMO. (**Figure 1C**). In parallel, brain KYNA levels in HET-*Kmo^+/−^* female and male mice were substantially elevated compared to WT-Control (**Figure 1D**). Together, these data demonstrate alteration in brain KP metabolism in both sexes of *Kmo* heterozygous mice.

### Disrupted sleep duration, architecture, and EEG power spectra in *Kmo* heterozygous mice

Elevated levels of brain KYNA have been shown to negatively impact sleep duration and architecture [33, 44, 66, 67]. Presently, we evaluated if sleep-wake behavior is disrupted in HET-*Kmo^+/−^* mice that we found have an endogenous elevation in brain KYNA due to disrupted KMO activity. During light phase, both sexes of HET-*Kmo^+/−^* mice spent less time in NREM sleep and more time awake compared to WT-Control mice (**Figure 2A**). Sex-specific differences were found when we evaluated sleep-wake architecture. Female HET-*Kmo^+/−^* mice had more wake bouts, while the bout number remained consistent between HET-*Kmo^+/−^* and WT-Control males for each vigilance state (**Figure 2B**). Moreover, during light phase HET-*Kmo^+/−^* females had shorter average NREM bout duration, while in HET-*Kmo^+/−^* male mice average REM bout duration was significantly reduced compared to their counterpart controls (**Figure 2C**). Sleep-wake behavior was also evaluated during the dark phase (**Supplementary Table 1**), and we determined that the average REM bout duration was significantly reduced in HET-*Kmo^+/−^* females compared to controls.

**Figure 2.**
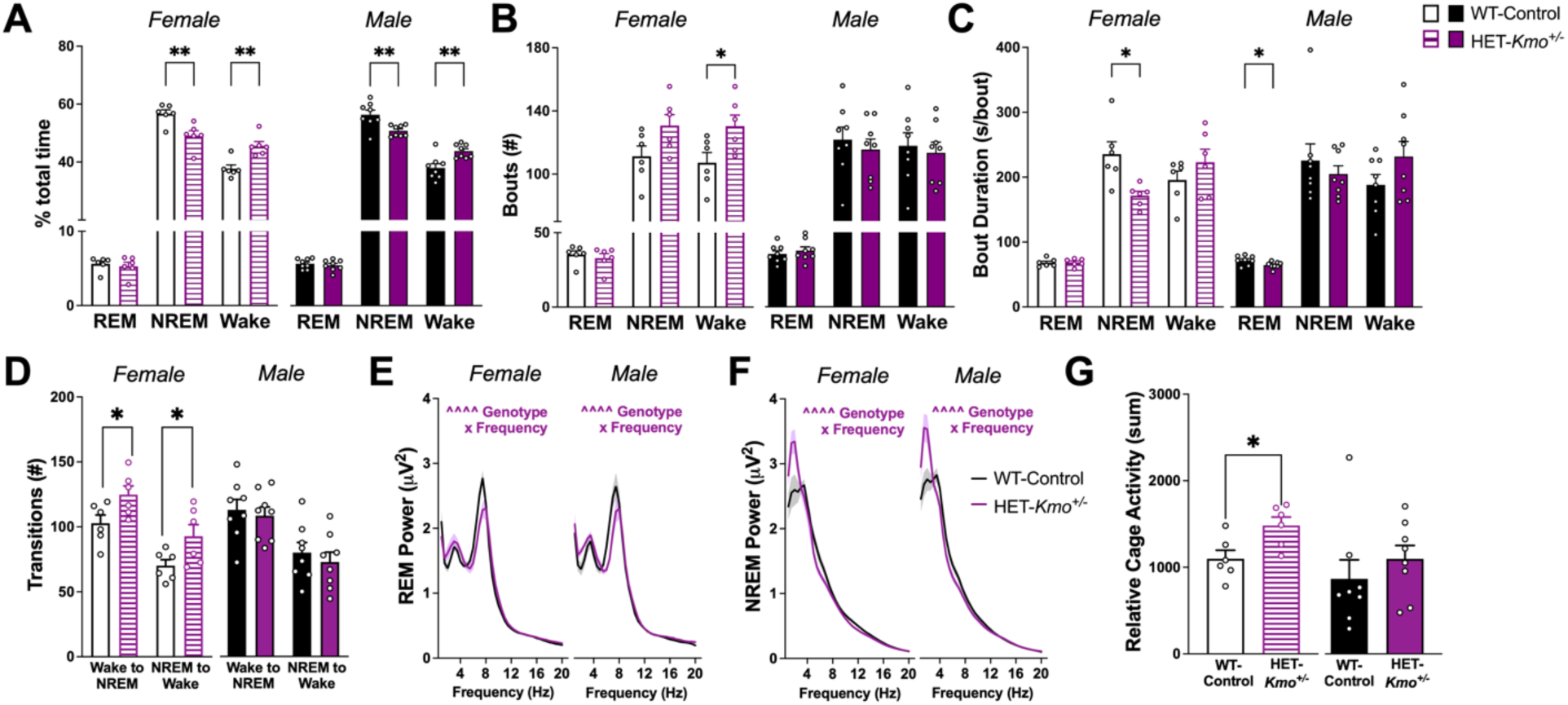
HET-*Kmo^+/−^* mice have decreased NREM sleep and more wake in comparison to wild-type (WT-Control) mice during the light phase. **(A)** Percentage of total time spent in each vigilance state (Unpaired t test **P<0.01). **(B)** Total number of bouts for each vigilance state (Unpaired t test *P<0.05). **(C)** Average bout duration for each vigilance state (Unpaired t test *P<0.05). **(D)** Number of vigilance state transitions (Unpaired t test *P<0.05). **(E)** REM sleep spectral power (Females: Two-way RM ANOVA Genotype x Frequency interaction F_(38, 380)_= 3.063, ^^^^P<0.0001; Males: Two-way RM ANOVA Genotype x Frequency interaction F_(38, 532)_= 3.109, ^^^^P<0.0001). **(F)** NREM sleep spectral power (Females: Two-way RM ANOVA Genotype x Frequency interaction F_(38, 380)_= 3.799, ^^^^P<0.0001; Males: Two-way RM ANOVA Genotype x Frequency interaction F_(38, 532)_= 5.967, ^^^^P<0.0001). **(G)** Relative cage activity (Unpaired t test *P<0.05). Data are mean ± SEM. N = 6-8 per group.

Sleep-wake architecture in HET-*Kmo^+/−^* and WT-Control mice was characterized by evaluating sleep onset latency and number of transitions between vigilance states. In male HET-*Kmo^+/−^* mice, latency to enter NREM sleep during light phase was significantly longer in comparison to their counterpart controls (**Supplementary Table 2**). Female HET-*Kmo^+/−^* mice transitioned more frequently from NREM sleep to wake and conversely from wake to NREM sleep compared to WT-Control females during the light phase, while these vigilance state transitions were unaltered in male mice (**Figure 2D**). All transitions for light and dark phase are presented in **Supplementary Table 2**.

Spectral power during NREM delta and REM theta were evaluated between HET-*Kmo^+/−^* and WT-Control mice of both sexes. During light phase, in both sexes, REM theta power was reduced in HET-*Kmo^+/−^* mice (**Figure 2E**), yet NREM delta power was significantly enhanced (**Figure 2F**). The sleep state power spectra differences persisted in dark phase, as shown in **Supplementary Figure 1**.

Increased wake duration in HET-*Kmo^+/−^* mice prompted an evaluation of home cage activity during the light phase. Female HET-*Kmo^+/−^* mice were significantly more active in the home cage in comparison to their counterpart controls, but no difference between genotypes were determined in male mice (**Figure 2G**).

### Mitochondrial respiration is altered in the hippocampal tissue of female *Kmo* heterozygous mice

KMO is localized to the outer mitochondrial membrane [51] and mitochondrial metabolic function is regulated by sleep and wake experiences [50, 68, 69]. Building on our finding that HET-*Kmo^+/−^* female mice spent less time in NREM sleep and more time awake compared to WT-Control counterparts, we next assessed mitochondrial respiration in the cortex and hippocampus, regions that are impacted by sleep quality, to evaluate brain region-specific respiratory rates.

Cortical mitochondrial complex I-linked oxygen consumption rate (OCR) was comparable between the two genotypes across all measured states: basal respiration, adenosine diphosphate (ADP) stimulated respiration (State 3), oligomycin inhibited adenosine triphosphate (ATP) synthase respiration (State 4o), and uncoupled respiration (State 3u) (**Figure 3A**). In contrast to cortical respiration, we observed a decrease in hippocampal complex II-linked State 3 respiration in HET-*Kmo^+/−^* females compared to WT-Control female mice (**Figure 3B**). Examination of cortical mitochondrial complex II-driven OCR revealed no significant differences in respiratory states between females of the two groups (**Figure 3C**). Both basal respiration and ADP stimulated respiration were significantly elevated in hippocampal complex II-dependent respiration in HET-*Kmo^+/−^* compared to WT-Control females (**Figure 3D**).

**Figure 3.**
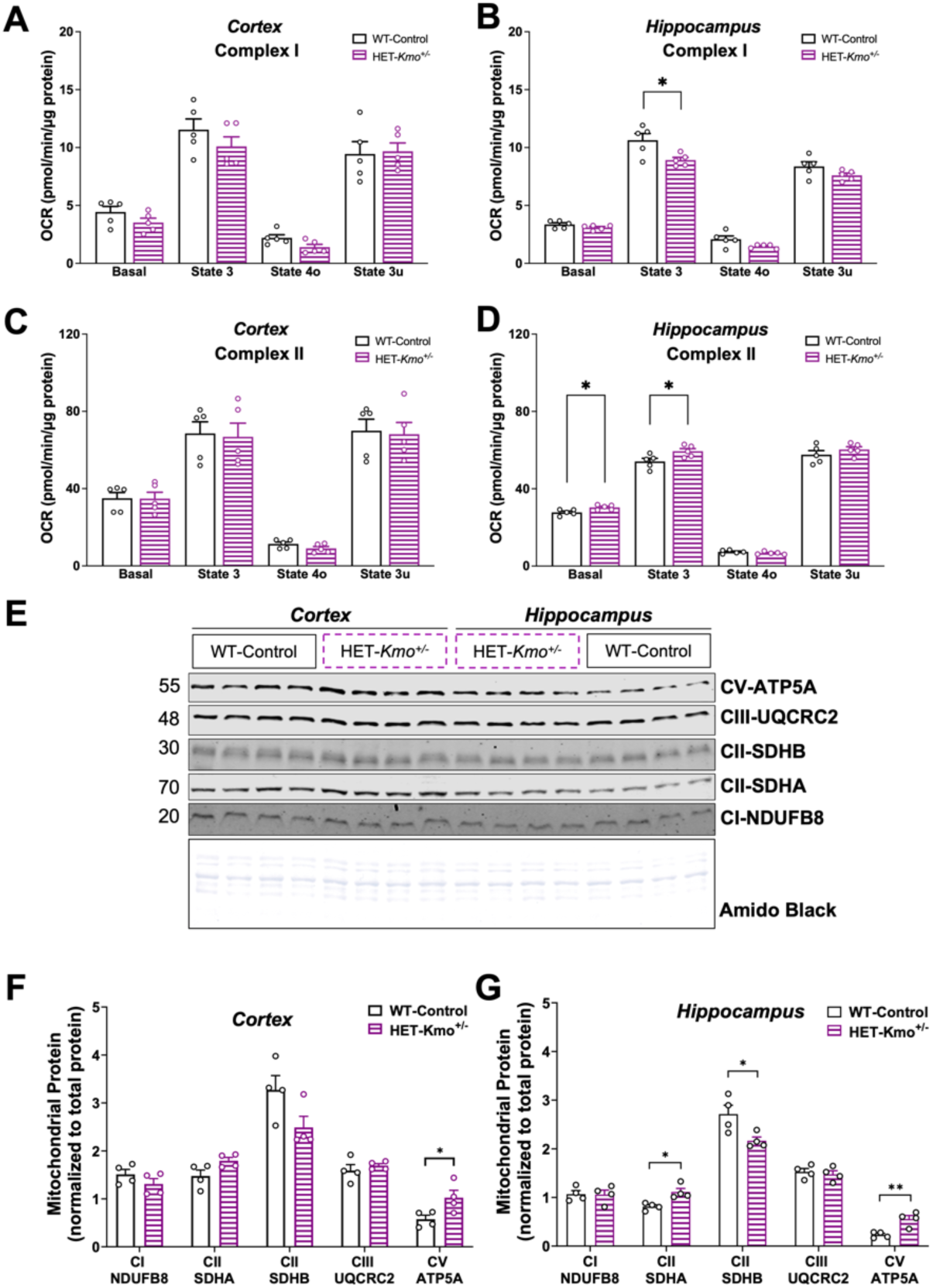
HET-*Kmo^+/−^* females display substrate-specific changes in hippocampal mitochondrial respiration. **(A)** Cortex complex I-dependent (glutamate/malate) respiration was unchanged between genotypes. **(B)** Hippocampus mitochondrial complex I-dependent State 3 (+ADP) respiration was significantly decreased in female HET-*Kmo^+/−^* versus WT-Control (Welch’s t test *P<0.05). **(C)** Cortex complex II-dependent (succinate/rotenone) respiration was unchanged between genotypes. **(D)** Hippocampus mitochondrial complex II-dependent respiration was significantly increased in both basal and State 3 respiration in female HET-*Kmo^+/−^* versus WT-Control (Welch’s t test *P<0.05). **(E)** Profiling of select mitochondrial oxidative phosphorylation subunits in isolated mitochondria demonstrated a significant upregulation of the ATP5A component of Complex V (CV) in both the cortex and hippocampus of HET-*Kmo^+/−^* female mice versus WT-Control. HET-*Kmo^+/−^* hippocampal mitochondria also showed a significant decrease in Complex II (CII) SDHB subunit, whereas the CII SDHA subunit increased significantly. **(F)** Quantification of mitochondrial proteins measured in cortex (Welch’s t test *P<0.05). **(G)** Quantification of mitochondrial proteins measured in hippocampus (Welch’s t test *P<0.05, **P<0.01). Data are mean ± SEM. N = 5 per group for respiratory data, N=4 per group for immunoblot analysis. Mitochondrial subunit levels were normalized to total protein load as determined by Amido Black staining.

Given brain region-dependent alterations in mitochondrial respiration, we sought to assess protein levels of mitochondrial electron transport chain (ETC) constituents, complex I-V, in the same cortical and hippocampal mitochondrial fractions (**Figure 3E**). We identified significantly increased protein levels of the cortical mitochondrial ETC Complex V ATP synthase F1 alpha (ATP5A) subunit in mitochondrial fractions from HET-*Kmo^+/−^* compared to WT-Control female mice (**Figure 3F**). The protein levels of the hippocampal mitochondrial fractions further emphasized the differences between WT-Control and HET-*Kmo^+/−^* groups. Specifically, evaluation of Complex II components revealed increased levels of the succinate dehydrogenase A (SDHA) subunit concomitant with decreased levels of the succinate dehydrogenase B (SDHB) subunit, while Complex V ATP5A subunit protein levels were significantly elevated in the hippocampus similar to the cortex of HET-*Kmo^+/−^* females (**Figure 3G**). In contrast, the levels of the Complex I subunit NADH:ubiquinone oxidoreductase B8 (NDUFB8) and the Complex III subunit ubiquinol-cytochrome c reductase core protein 2 (UQCRC2) were comparable between the two genotypes in both cortical and hippocampal mitochondrial fractions.

In addition to ATP production, OXPHOS generates reactive oxygen species (ROS) as a derivative [70]. Overproduction of ROS leads to oxidative stress, which is implicated in the pathophysiology of several conditions, including sleep-wake homeostasis [71–73]. Oxidative stress can be determined by measuring protein adducts formed by 4-hydroxy-2-nonenal (HNE), a product of lipid peroxidation [74]. We examined HNE-modified protein adducts in both the cortical and hippocampal tissue from WT-Control and HET-*Kmo^+/−^* female mice, and we found no significant differences in HNE adducts between the two genotypes (**Supplementary Figure 2B, 2C**). KMO enzyme catalyzes hydroxylation of kynurenine to 3-HK. Autoxidation of 3-HK has been shown to promote ROS production, and consequently, oxidative stress [75, 76]. We examined 3-HK levels in the brain tissue of HET-*Kmo^+/−^* compared to WT-Control females. As expected, due to the reduced KMO enzyme activity, levels of 3-HK in the brain were significantly decreased in HET-*Kmo^+/−^* female mice relative to control counterparts (**Supplementary Figure 2D**). Since the *Kmo* genotype directs the KMO enzyme protein levels on the outer mitochondrial membrane, we further evaluated protein levels of another abundant outer mitochondrial membrane marker, the voltage-dependent anion channel (VDAC) 2. No differences in VDAC2 protein levels were detected between WT-Control and HET-*Kmo^+/−^* groups in either cortical or hippocampal mitochondria (**Supplementary Figure 2E**)

Taken together, these data suggest that *Kmo* heterozygous females predominantly exhibit substrate-specific changes in hippocampal mitochondrial respiration, concurrent with compensatory changes in the mitochondrial ETC subunits protein levels, without apparent oxidative stress that may in part be due to reduced 3-HK brain levels.

### Elevated kynurenine levels in the breast milk of *Kmo* heterozygous mothers

Kynurenine and KYNA levels in plasma and placenta of HET-*Kmo^+/−^* mothers have previously been reported [21] to be comparable to WT-Control mothers, pointing to similar *in utero* exposure to the KP metabolites during offspring embryonic development. To further elucidate the postnatal effect of maternal genotype on developing offspring, we presently assessed the KP metabolites in breast milk of both WT-Control and HET-*Kmo^+/−^* mothers. Breast milk tryptophan levels did not differ between mothers of the two genotypes; however, kynurenine levels were significantly higher (3-fold) in HET-*Kmo^+/−^* mothers compared to controls generating over 3-fold increased kynurenine to tryptophan ratio in HET-*Kmo^+/−^* mothers (**Figure 4A, 4B**). Notably, breast milk corticosterone levels were proportionate between HET-*Kmo^+/−^* mothers and their counterpart controls (**Figure 4C**). Breast milk from *Kmo* heterozygous mothers may thereby directly deliver elevated kynurenine to offspring during early postnatal development.

**Figure 4.**
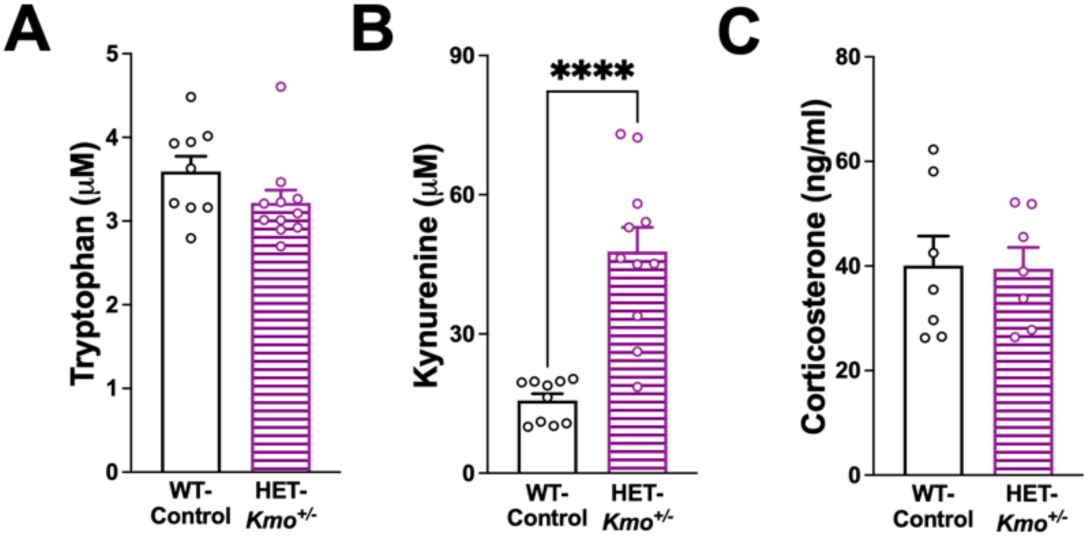
Breast milk content of kynurenine is elevated in HET-*Kmo^+/−^* compared to WT-Control mothers. **(A)** Maternal breast milk tryptophan. **(B)** Maternal breast milk kynurenine (Unpaired t test ****P<0.0001). **(C)** Maternal breast milk corticosterone. Data are mean ± SEM. N = 7-11 per group.

Of note, no major differences in parental care behaviors were observed between HET-*Kmo^+/−^* and WT-Control parents in three studied categories: positive behaviors (grooming pups, contact with pups, nursing), self-care (eating and drinking, self-grooming), and negative behaviors (nest maintenance, parent out of nest) (**Supplementary Figure 3**, **Supplementary Table 3**).

### Learning impairments in wild-type female offspring from *Kmo* heterozygous parents

Behavioral studies were conducted in adult wild-type (WT) offspring from two distinct breeding groups: i) WT-Control parents generated WT-Control offspring and ii) *Kmo* heterozygous (HET-*Kmo^+/−^* x HET-*Kmo^+/−^*) parents generated WT-*Kmo^+/+^* offspring (**Figure 5A**). Of note, in the latter breeding pair ~25% of offspring were WT-*Kmo^+/+^*.

**Figure 5.**
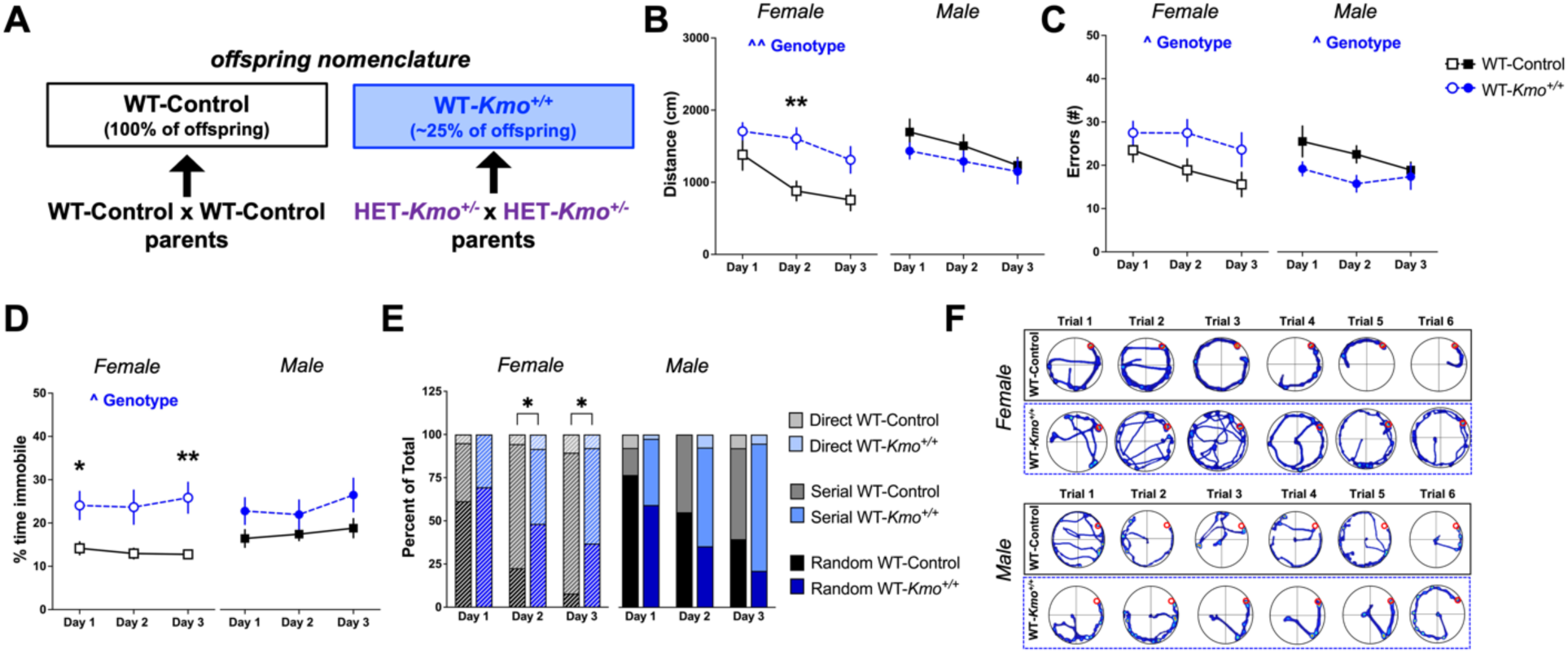
Impaired spatial learning in female wild-type (WT-*Kmo^+/+^*) offspring from HET-*Kmo^+/−^* parents. **(A)** Schematic representation of the experimental offspring groups. WT-Control offspring are derived from WT-Control parents. WT-*Kmo^+/+^* offspring are derived from heterozygous *Kmo* (HET-*Kmo^+/−^*) parents. **(B)** Distance traveled in the Barnes maze (Female: Two-way RM ANOVA Genotype effect F_(1, 36)_= 11.08, ^^P<0.01 with Bonferroni’s post hoc test **P<0.01). **(C)** Errors in the Barnes maze (Female: Two-way RM ANOVA Genotype effect F_(1, 36)_= 5.311, ^P<0.05; Male: Two-way RM ANOVA Genotype effect F_(1, 40)_= 4.391, ^P<0.05). **(D)** Percent of time immobile in the Barnes maze (Female: Two-way RM ANOVA Genotype effect F_(1, 24)_= 7.1, ^P<0.05 with Bonferroni’s post hoc test *P<0.05, **P<0.01). **(E)** Percent of time spent in each search strategy on the Barnes maze (Chi-square distribution test *P<0.05). **(F)** Representative images of video tracking in the Barnes maze across learning trials. Data are mean ± SEM. N = 19-21 per group.

Barnes maze testing was employed to evaluate learning and memory. Distance traveled, an indicator of learning across acquisition days, was significantly greater in WT-*Kmo^+/+^* female mice in comparison with WT-Control (main effect of genotype), particularly on the second learning day. This finding is unlikely due to hyperactivity since the average velocity across learning days was not significantly different between females of the two genotypes (data not shown). Male WT-*Kmo^+/+^* mice moved similar distance to their counterpart controls (**Figure 5B**). Compared to controls, WT-*Kmo^+/+^* female mice committed significantly more errors, coinciding with longer distance traveled and an overall indication of poorer learning in the Barnes maze. Conversely, WT-*Kmo^+/+^* males committed fewer errors across days in the task than WT-Control mice (**Figure 5C**). Consequently, latency to entering the escape box was significantly impacted by genotype in female offspring only (**Supplementary Figure 4A**). We also noted that during the learning days, WT-*Kmo^+/+^* female, but not male, mice were spending significantly more time immobile than their counterpart controls (main effect of genotype) (**Figure 5D**). Taken together, longer distance traveled, increased errors, and increased immobility in WT-*Kmo^+/+^* females pinpoint poor spatial learning in the Barnes maze.

We evaluated three different search strategies employed by mice to locate the Barnes maze escape box: random, serial, or direct. WT-*Kmo^+/+^* females continued to utilize the less efficient random search strategy to the greater extent than WT-Control females on the second and the third learning days. No differences in utilization of different search strategies were determined between male WT-*Kmo^+/+^* and their counterpart controls (**Figure 5E**). Representative traces of movement on the Barnes maze demonstrate stark differences between sexes (**Figure 5F**). Altogether, Barnes maze data collectively indicate impaired spatial learning and memory formation in female, but not male WT-*Kmo^+/+^* offspring compared to controls.

### Impaired reversal learning in male wild-type offspring from *Kmo* heterozygous parents

We next tested reversal learning in the Barnes maze to evaluate cognitive flexibility mediated by the prefrontal cortex [77, 78]. While no significant impairments in reversal learning were found in female mice, male WT-*Kmo^+/+^* mice traveled significantly longer distance (**Figure 6A**) and committed more errors (**Figure 6B**) when compared to male WT-Control mice. No differences in the distribution of search strategies employed during the reversal trial were found for both sexes (**Figure 6C**), as indicated by the representative track plots (**Figure 6D**). Lastly, time spent immobile on the maze during the reversal trial did not differ between female and male mice of both genotypes (**Supplementary Figure 4B**). Taken together, reversal trial Barnes maze revealed that male, but not female WT-*Kmo^+/+^* offspring have impairments in cognitive flexibility in a spatial navigation task.

**Figure 6.**
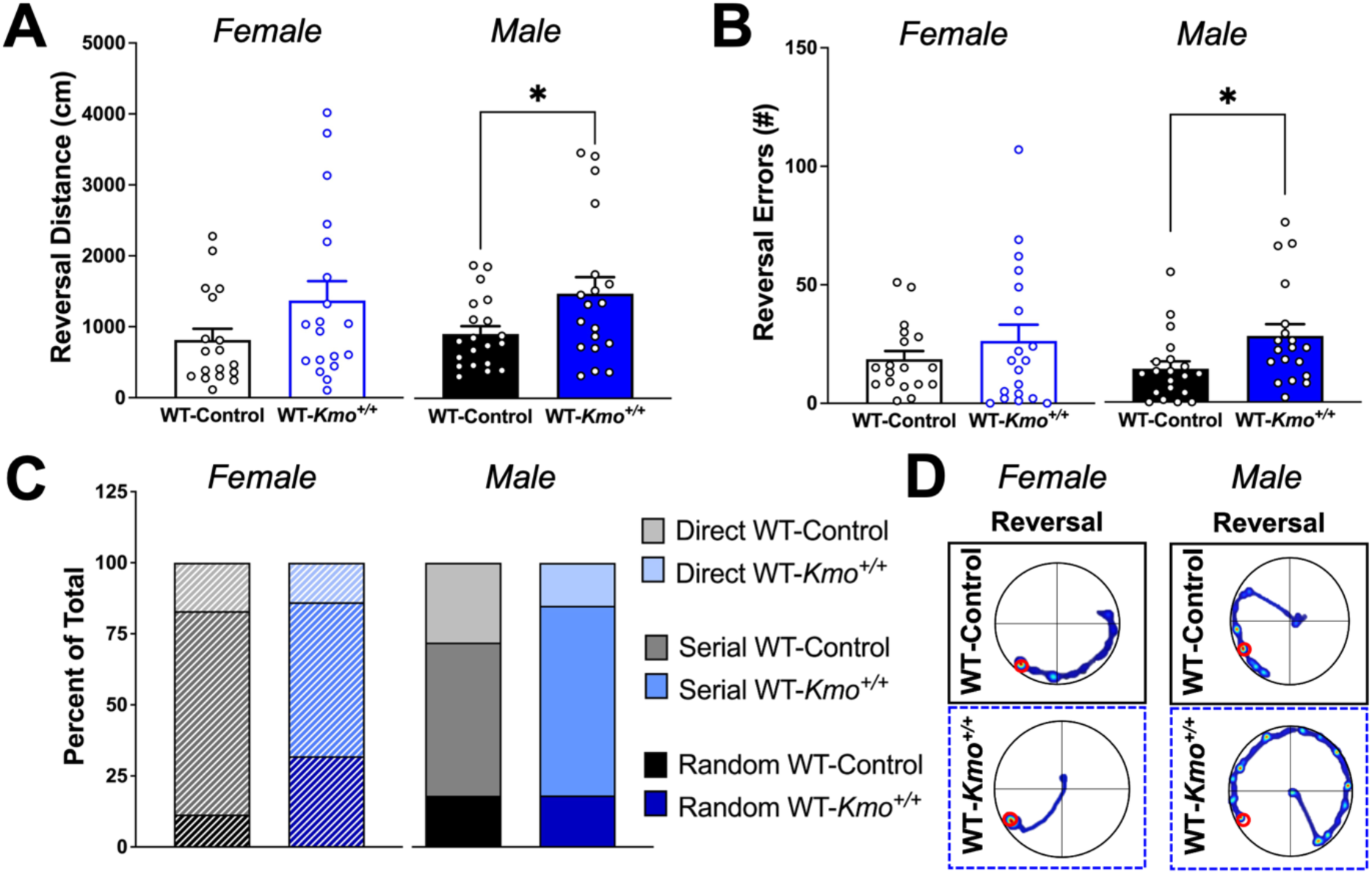
Impaired reversal learning in male wild-type (WT-*Kmo^+/+^*) offspring from HET-*Kmo^+/−^* parents. **(A)** Distance traveled in the reversal trial of the Barnes maze (Unpaired t-test *P<0.05). **(B)** Errors in the reversal trial of the Barnes maze (Unpaired t-test *P<0.05). **(C)** Percent of time spent in each search strategy in the reversal trial of the Barnes maze. **(D)** Representative images of video tracking in the Barnes maze during the reversal trial. Data are mean ± SEM. N = 19-21 per group.

### Wild-type offspring from *Kmo* heterozygous parents display decreased amount of exploration

To further understand enhanced immobility in the Barnes maze paradigm, we employed testing with the elevated zero maze to better understand exploratory and anxiety-like behavior in adult offspring. Both male and female WT offspring (WT-*Kmo^+/+^*) derived from *Kmo* heterozygous parents, traveled significantly less distance in the elevated zero maze (**Figure 7A**). Female WT-*Kmo^+/+^* mice entered both open and closed areas less frequently compared to controls (WT-Control), but all male mice entered maze arms at the same frequency (**Figure 7B**).

**Figure 7.**
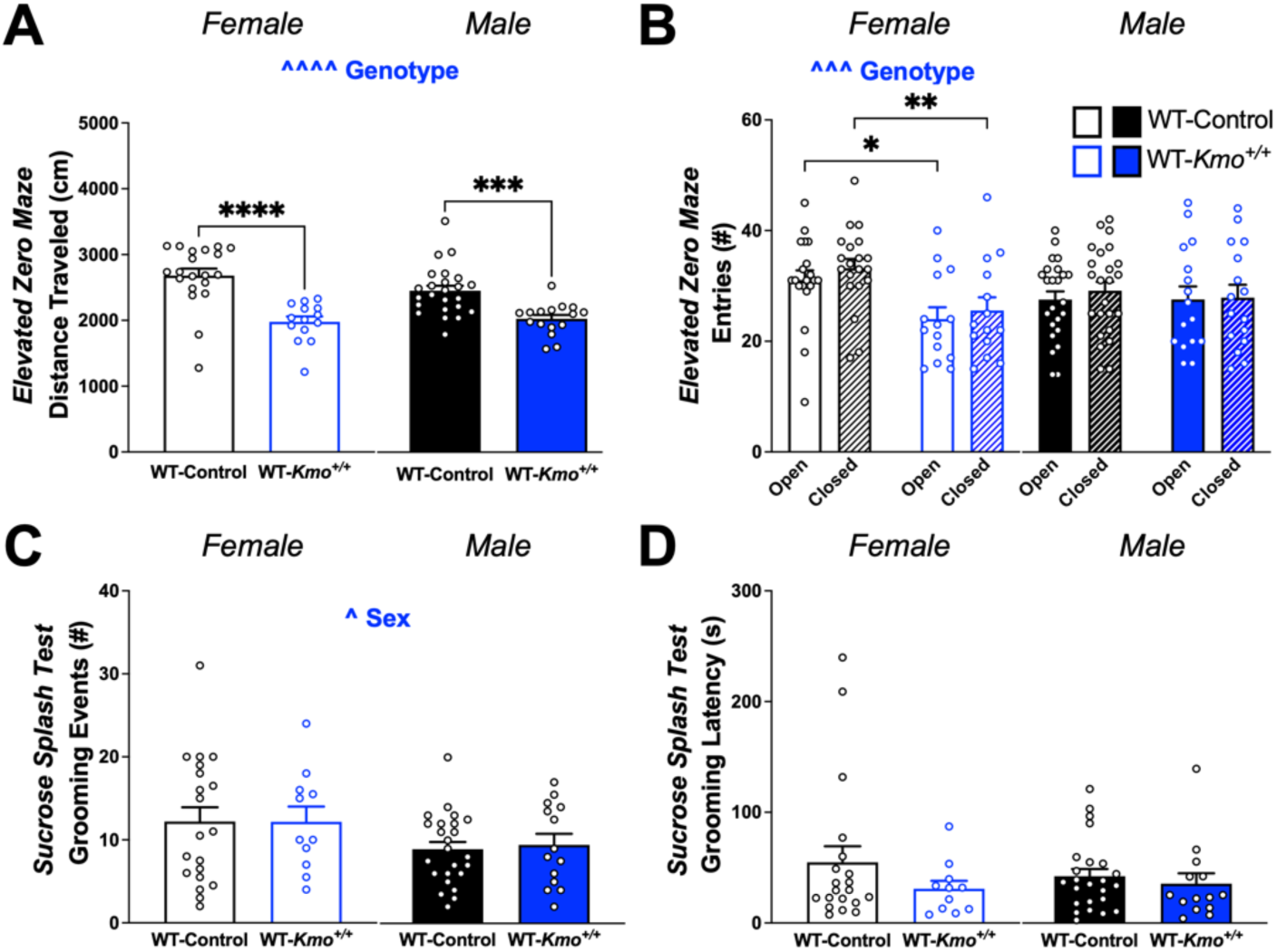
Increased anxiety-like behavior in female wild-type (WT-*Kmo^+/+^*) offspring from HET-*Kmo^+/−^* parents. **(A)** Distance traveled in the elevated zero maze (Two-way RM ANOVA Genotype effect F_(1, 70)_= 44.25, ^^^^P<0.0001 with Fisher’s LSD post hoc test ***P<0.001, ****P<0.0001). **(B)** Entries to open and closed area in the elevated zero maze (Females: Two-way RM ANOVA Genotype effect F_(1, 64)_= 14.71, ^^^P<0.001 with Fisher’s LSD post hoc test *P<0.05, **P<0.01). **(C)** Number of grooming events in the sucrose splash test (Two-way RM ANOVA Sex effect F_(1, 65)_= 4.093, ^P<0.05). **(D)** Latency to grooming in the sucrose splash test. Data are mean ± SEM. N = 11-24 per group.

To evaluate if reduced exploration in WT-*Kmo^+/+^* mice was related to a lack of motivation [77], we performed a sucrose splash test to assess motivation of self-care (grooming) behavior in adult offspring. The number of groom events was significantly lower in male compared to female mice (**Figure 7C**), but we did not detect any change in grooming latency or differences related to parental genotype (**Figure 7D**).

### Reduced NREM sleep and increased wakefulness in wild-type female offspring from *Kmo* heterozygous parents

The finding of sex-specific learning impairments among offspring from HET-*Kmo^+/−^* parents led us to investigate sleep behavior in a separate cohort of female and male WT-*Kmo^+/+^* mice compared to counterpart controls, as sleep significantly contributes to short- and long-term memory formation and consolidation [79–81].

Total duration of NREM sleep was reduced across the light phase in WT-*Kmo^+/+^* female mice compared to WT-Control, and consequently, wakefulness was significantly increased between ZT 4 and ZT 12. Statistical analyses point to a significant effect of genotype on wake duration and an approaching trend for NREM duration. No changes in NREM sleep and wake duration were found in male offspring during light phase (**Figure 8A, 8B**). While total sleep duration was significantly reduced in female WT-*Kmo^+/+^* mice, the total REM duration was not impacted by parental genotype of offspring in both sexes during light phase (**Supplementary Table 4**).

**Figure 8.**
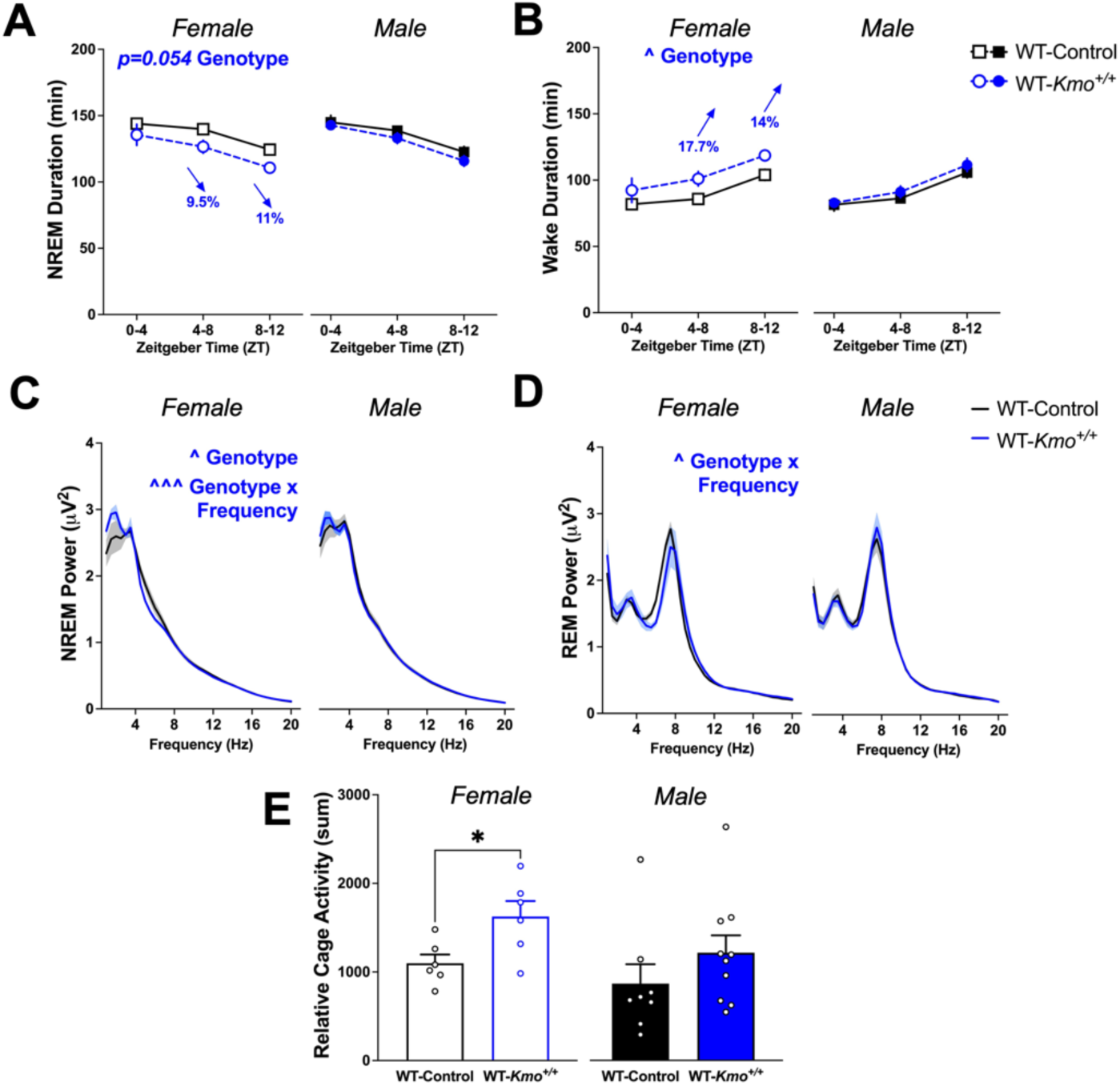
Female wild-type (WT-*Kmo^+/+^*) from HET-*Kmo^+/−^* parents have reduced NREM sleep and more wake in comparison to wild-type (WT-Control) mice during the light phase. **(A)** Total NREM duration (Females: Two-way RM ANOVA Genotype effect F_(1, 10)_= 4.776, P=0.054). **(B)** Total wake duration (Females: Two-way RM ANOVA Genotype effect F_(1, 10)_= 5.584, ^P<0.05). **(C)** NREM sleep spectral power (Females: Two-way RM ANOVA Genotype effect F_(1, 10)_= 8.711, ^P<0.05, Genotype x Frequency interaction F_(38, 380)_= 2.022, ^^^P<0.001). **(D)** REM sleep spectral power (Females: Two-way RM ANOVA Genotype x Frequency interaction F_(38, 380)_= 1.487, ^P<0.05). **(E)** Relative cage activity (Unpaired t test: *P<0.05). Data are mean ± SEM. N = 6-10 per group.

Spectral analysis of sleep EEG revealed sex-specific modifications in low-frequency delta oscillation bands (0.5-4 Hz) during light phase. A main effect of genotype as well as a genotype x frequency interaction for NREM power was determined in female offspring. Delta spectral power in WT-*Kmo^+/+^* females was enhanced compared to WT-Control female mice. No differences in NREM delta spectra were detected among both groups of male offspring, WT-*Kmo^+/+^* and WT-Control (**Figure 8C**). Further assessment of low frequency theta oscillation bands (4-8 Hz) indicated sex-dependent changes in REM power during light phase. In comparison to their counterpart controls, female WT-*Kmo^+/+^* offspring displayed reduced theta spectral power noted as a significant interaction between genotype and frequency. We observed no differences in REM theta spectra between WT-*Kmo^+/+^* and WT-Control male mice (**Figure 8D**).

Wake behavior was assessed by analysis of relative cage activity. Since female WT-*Kmo^+/+^* offspring exhibited prolonged wakefulness during light phase, expectedly, we noticed an increase in cage activity compared to their counterpart controls. Relative activity of WT-*Kmo^+/+^* and WT-Control male offspring did not differ between groups (**Figure 8E**).

Number of bouts and average bout duration during REM sleep, NREM sleep, and wake were evaluated to further understand vigilance state architecture (**Supplementary Table 4**). Of note, average wake bout duration was significantly impacted by genotype in male offspring during light phase. EEG spectra analysis within the delta and theta ranges revealed sex-dependent differences during the dark phase as well. An interaction between genotype and frequency was found in female WT-*Kmo^+/+^* mice, such that NREM delta power was increased compared to WT-Control females (**Supplementary Figure 5A**). In male WT-*Kmo^+/+^* mice we observed a main effect of genotype and enhanced REM theta power (**Supplementary Figure 5B**). Together, these data imply noteworthy disturbances in sleep-wake behavior in only female, but not male, WT-*Kmo^+/+^* offspring compared to counterpart control offspring.

## DISCUSSION

Clinical studies have shown that both the genes parents pass on to their children and the genes they do not transmit can influence traits. This occurs not only through direct inheritance but also by shaping the environment and nurturing behaviors that impact a child’s development [82, 83]. Preclinical studies, such as the *Kmo* heterozygous model used in this study, are crucial for disentangling how parental genetic and environmental factors independently and interactively influence offspring development [84]. Here, we report that parental *Kmo* genotype affects the development of sex-specific cognitive and behavioral traits in offspring through genetic nurturing effects.

Inheritable genetic factors play critical roles in brain development by regulating synaptic plasticity and neural circuitry, and many are implicated in neurodevelopmental disorders [85–89]. The KMO model used here was designed to investigate the KP’s involvement in neuropsychiatric endophenotypes, including related biochemical and neurobehavioral abnormalities [12, 21, 53]. While *Kmo* knockout mice provide insights into complete loss-of-function effects, such as reduced 3-HK levels, elevated KYNA, memory deficits, and anxiety-like behaviors [12, 90] - complete *Kmo* loss is not observed in humans. Instead, we focused on *Kmo* heterozygous mice to study partial *Kmo* loss and its genetic nurturing effects on wild-type offspring. We identified parental phenotypes in *Kmo* heterozygous mice likely to contribute to offspring neurodevelopmental outcomes (**Figure 9**).

**Figure 9.**
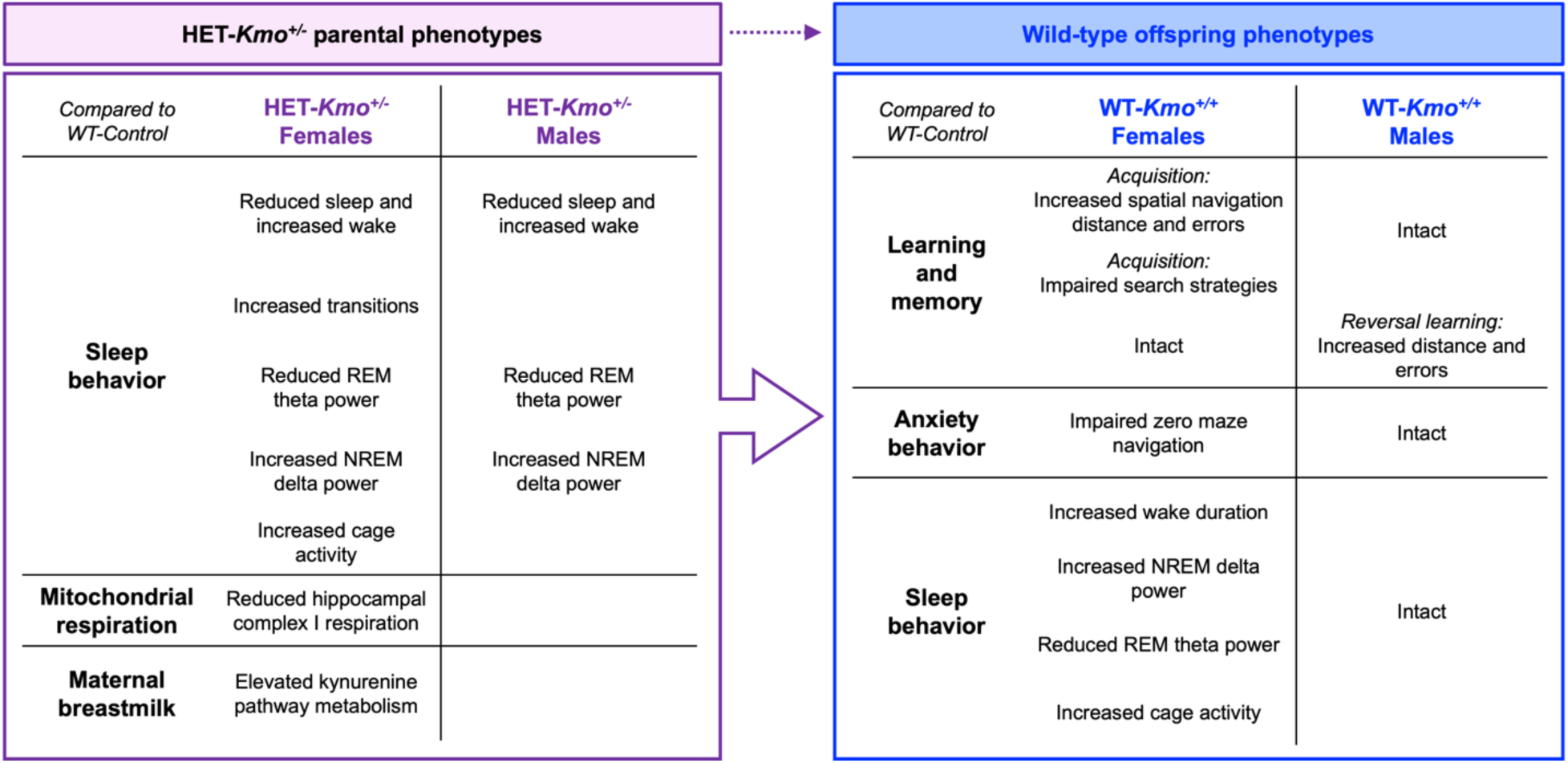
Schematic representation of major findings supporting the conclusion that parental HET-*Kmo^+/−^* genotype influences wild-type offspring phenotypes. Disruptions in sleep were observed in female and male HET-*Kmo^+/−^* mice, and HET-*Kmo^+/−^* female mice had altered hippocampal mitochondria respiration and increased kynurenine levels in breast milk. Even though WT-*Kmo^+/+^* inherit identical genetic material as WT-Control, the present data suggest that behavioral phenotypes of offspring could be shaped by parental physiology, behavior, or metabolic state influenced by their *Kmo* genotype. Sex-specific differences are noted in offspring. Female WT-*Kmo^+/+^* offspring from HET-*Kmo^+/−^* parents exhibit spatial learning impairments, increased anxiety-like behavior, and sleep disturbances. Male WT-*Kmo^+/+^* offspring from HET-*Kmo^+/−^* parents display impaired reversal learning, but intact spatial learning, anxiety-like behavior, and sleep patterns.

Biochemical analysis determined that in *Kmo* heterozygous mice, brain KMO activity was reduced, and KYNA levels were increased compared to controls. These findings align with postmortem analysis showing elevated KYNA levels and reduced KMO in the brains of individuals with schizophrenia [17, 27, 28]. Elevated KYNA levels have also been observed in cerebrospinal fluid (CSF) levels of those with schizophrenia and bipolar disorder [91–93], correlating with cognitive impairments including executive function [94]. In preclinical models, elevated KYNA impairs spatial learning, memory, and cognitive flexibility, likely through modulation of cholinergic and glutamatergic neurotransmission [34, 35, 95, 96].

The sleep deficiencies identified in HET-*Kmo*^+/−^ mice support growing literature that increased KYNA reduces sleep duration and impairs sleep architecture [33, 44, 66, 67]. Importantly, our study extends these findings by demonstrating that elevated KYNA and reduced KMO activity in HET-*Kmo*^+/−^ mice disrupt sleep behavior and mitochondrial function, processes closely tied to neurodevelopment. Sleep disruptions included shorter NREM sleep duration, prolonged wakefulness, and fragmented sleep architecture, especially in females. The increased low-frequency NREM sleep delta oscillations suggest a compensatory need for sleep that persisted across light phases in *Kmo* heterozygous female mice.

Sleep disruptions in female HET-*Kmo^+/−^* mice prompted further investigation into the interplay between sleep and mitochondrial function, processes tightly regulated by respiration to optimize cellular function [47–49]. The brain’s energy requirements vary by vigilance state, with higher demands during wakefulness and reduced needs during sleep [97, 98]. ATP generated by mitochondria through oxidative phosphorylation (OXPHOS) is specifically adapted to meet these changing energy demands [99]. In *Drosophila*, loss of the KMO homologue reduced the respiratory capacity of mitochondrial electron transport chain (ETC) complex I and elevated mitochondrial mass [51]. Extending these findings, we observed decreased complex I-driven respiration and increased complex II-dependent respiration in hippocampal mitochondria of *Kmo* heterozygous females, suggesting enhanced complex II activity compensates for elevated energy demands during prolonged wakefulness. This was mirrored by alterations in hippocampal the Complex II subunits SDHA and SDHB, suggesting adaptive remodeling of Complex II. Changes in mitochondrial protein levels may further affect cortico-hippocampal synaptic plasticity, linking mitochondrial function to neurotransmission and plasticity [100]. It is also noteworthy that increased levels of ATP5A, a component of the catalytic domain of ATP synthase (complex V) were increased in the *Kmo* heterozygotes vs. WT controls. Future studies should explore region-specific mitochondrial alterations and their effects on cognition and memory in *Kmo* heterozygous mice.

To further explore the impact of maternal *Kmo* heterozygosity on offspring outcomes through genetic nurture, we studied breast milk composition. This study is the first to quantify KP metabolites in the breast milk of lactating mice, the primary source of nutrition for offspring during the critical early postnatal weeks of development. While KP metabolite levels in plasma and placenta were similar between *Kmo* heterozygous and control mothers [21], *Kmo* heterozygous mothers had elevated kynurenine levels in breast milk, which could promote increased neosynthesis of KYNA and 3-HK in offspring during a developmental window critical for neurogenesis, synaptogenesis, and myelination [101]. Despite the observed biochemical and sleep behavioral changes in *Kmo* heterozygous parents, parental care behaviors were similar across genotypes, indicating that early postnatal development was not significantly influenced by differences in caregiving [102, 103]. These findings suggest that changes in offspring outcomes may primarily arise from biochemical factors rather than variations in parental care. However, future studies utilizing cross-fostering designs would provide more conclusive evidence by disentangling the effects of maternal *Kmo* genotype from postnatal environmental influences on offspring development.

Wild-type offspring from *Kmo* heterozygous parents exhibited behavioral sex-dependent differences. Female wild-type offspring from *Kmo* heterozygous parents displayed pronounced spatial learning impairments and heightened anxiety-like behavior. These impairments were characterized by increase distance, more errors, and a persistent reliance on the less effective random search strategy in the Barnes maze. Interestingly, female wild-type offspring from *Kmo* heterozygous parents were more immobile during spatial navigation. This heightened immobility may stem from anxiety or reduced exploratory motivation, potentially hindering task performance. Females also exhibited reduced exploration in the elevated zero maze, yet despite these deficits, females did not display anhedonia or lack of motivation to groom during splash testing. Conversely, male wild-type offspring from *Kmo* heterozygous parents exhibited deficits only during the reversal trial in spatial navigation, which is associated with prefrontal cortex-mediated cognitive flexibility [104]. Sex differences in cognitive flexibility, and notable in the reversal trial of the Barnes maze, have been previously reported [45, 105].

Accompanying these cognitive impairments were sex-specific alterations in sleep patterns, highlighting a potential mechanistic role for sleep in cognitive outcomes. Female, but not male, wild-type offspring from *Kmo* heterozygous parents showed reduced NREM sleep duration and reduced theta power during REM sleep, accompanied by prolonged wakefulness. As both NREM and REM sleep are important for memory consolidation [81, 106], these disruptions likely contributed to the observed cognitive deficits. Reduced REM sleep theta power in female offspring further suggests impaired organization of spatial memory engrams [107, 108]. These findings align with literature demonstrating that NREM sleep modulates hippocampal-dependent learning via encoding and synaptic scaling [109, 110].

In contrast, wild-type male offspring from Kmo heterozygous parents displayed no significant disruptions in sleep behavior compared to controls, apart from the peculiar, elevated theta power during REM sleep in the dark phase. Elevated theta oscillations, which synchronize hippocampal-prefrontal activity, may support cognitive resilience by enhancing spatial memory retrieval and decision-making processes [111, 112]. However, context-specific theta activity during the Barnes maze, which we were unable to measure in the present paradigm, could potential also reflect compensatory mechanism that, while advantageous in certain scenarios, might compromise executive function during reversal learning [113].

Sleep plays a critical role in memory consolidation, and disruptions in sleep among individuals with neurodevelopmental disorders may exacerbate clinical symptoms, including cognitive challenges [109, 110]. Despite this, the interplay between sleep and cognitive mechanisms, particularly how they intersect with sex differences commonly observed in individuals with autism, ADHD, and schizophrenia, remains poorly understood. Furthermore, in rodent studies, sex differences in sleep patterns and the impact of sex hormones on sleep regulation have been well-documented. Estrogen has been shown to modulate sleep homeostasis in female rodents, promoting increased wakefulness during the dark period [114–117]. However, the mechanisms by which sex hormones and associated differences in sleep patterns contribute to cognitive impairments in neurodevelopmental disorders remain to be elucidated.

## CONCLUSIONS

The sex-specific phenotypes observed in offspring from *Kmo* heterozygous parents likely result from interactions between genetic, epigenetic, and environmental factors. Males may exhibit resilience due to intact NREM sleep architecture or enhanced prefrontal-hippocampal synchronization, which supports synaptic plasticity and memory. Additionally, differential parental contributions and inherited epigenetic modifications may further shape cognitive and sleep outcomes [118, 119].

Overall, the data underscore the critical interplay between sleep and cognitive function in offspring from *Kmo* heterozygous parents and suggest that sex-dependent mechanisms – potentially involving differences in brain network oscillations and parental influences – contribute to the observed phenotypic divergence. Further studies are warranted to elucidate the specific neural and molecular pathways underpinning male resilience and female vulnerability in the wild-type offspring from *Kmo* heterozygous parents. These findings contribute to a growing body of evidence highlighting the interplay between genetic and environmental factors in shaping offspring endophenotypes. By advancing our understanding of how parental *Kmo* genetics influence sleep, cognition, and behavior, this work may provide valuable insights into the etiology of sex-specific vulnerabilities observed in neurodevelopmental disorders, ultimately informing targeted interventions and therapeutics.

## DECLARATIONS

### Ethics approval and consent to participate

no applicable

### Consent for publication

All authors consent to publication of the work.

### Availability of data and materials

The data that support the findings of this study are available from the corresponding author upon reasonable request.

### Authors’ contributions

<u>Snezana Milosavljevic:</u> writing – original draft; writing – review and editing; visualization; investigation; conceptualization; formal analysis; methodology, funding acquisition. <u>Maria V. Piroli:</u> writing – review and editing; investigation; visualization; conceptualization; formal analysis; methodology. <u>Emma J. Sandago:</u> writing – review and editing; investigation; visualization; conceptualization; formal analysis; methodology. <u>Gerardo G. Piroli:</u> writing – review and editing; investigation; visualization; conceptualization; formal analysis; methodology. Holland H. Smith: writing – review and editing; investigation; visualization; methodology. <u>Sarah Beggiato</u>: Conceptualization; investigation; writing – review and editing; formal analysis. <u>Norma Frizzell</u>: Conceptualization; investigation; funding acquisition; writing – original draft; visualization; writing – review and editing; formal analysis; project administration; supervision; resources. <u>Ana Pocivavsek</u>: Conceptualization; investigation; funding acquisition; writing – original draft; visualization; writing – review and editing; formal analysis; project administration; supervision; resources.

## Supporting information

Supplemental Statistical Table

## Acknowledgements

The authors would like to thank Morgan Lambert and Sarisha Menon for their technical contributions to this work.

## Supplemental Materials

### Supplemental Statistical Table

All three-way and two-way ANOVA statistical results are presented in the supplemental .xls file named “*Milosavljevic et al_Supplemental Statistical Table*” to denote statistical analysis for manuscript figures and supplemental figures wherein male and female data are evaluated simultaneously.

**Supplementary Table 1.** Sleep-wake parameters in HET-*Kmo^+/−^* mice in comparison to wild-type (WT-Control) mice during the dark phase. Data are mean ± SEM. Unpaired t test: *P<0.05 vs. WT-Control. N = 6-8 per group.

| Dark Phase | Female |  | Male |  |
| --- | --- | --- | --- | --- |
|  | WT-Control | HET-*Kmo*<sup>+/-</sup> | WT-Control | HET-*Kmo*<sup>+/-</sup> |
| REM Duration (% total time) | 2.4 ± 0.1 | 2.2 ± 0.3 | 2.9 ± 0.2 | 2.8 ± 0.2 |
| NREM Duration (% total time) | 34 ± 3 | 31 ± 4 | 36 ± 2 | 34 ± 2 |
| Wake Duration (% total time) | 63 ± 3 | 66 ± 4 | 61 ± 2 | 64 ± 2 |
| REM Bouts (#) | 15 ± 1 | 15 ± 1 | 18 ± 1 | 20 ± 2 |
| NREM Bouts (#) | 71 ± 5 | 87 ± 12 | 92 ± 6 | 82 ± 6 |
| Wake Bouts (#) | 74 ± 5 | 90 ± 11 | 93 ± 6 | 85 ± 6 |
| REM Bout Duration (s/bout) | 77 ± 5 | **65 ± 2 \*** | 72 ± 3 | 64 ± 3 |
| NREM Bout Duration (s/bout) | 223 ± 24 | 167 ± 8 | 178 ± 12 | 187 ± 10 |
| Wake Bout Duration (s/bout) | 769 ± 92 | 908 ± 230 | 619 ± 127 | 631 ± 81 |
| Relative Cage Activity (sum) | 2964 ± 443 | 3304 ± 662 | 2227 ± 288 | 2512 ± 325 |

**Supplementary Table 2.** Sleep onset and number of transitions between vigilance states in female and male mice during light and dark phase. Data are mean ± SEM. Unpaired t test: *P<0.05 vs. WT-Control. N = 6-10 per group.

| Light Phase | Female |  |  |
| --- | --- | --- | --- |
|  | WT-Control | WT-*Kmo*<sup>+/+</sup> | HET-*Kmo*<sup>+/-</sup> |
| NREM Onset (min) | 4.9 ± 1.8 | 6.7 ± 1.2 | 12.2 ± 3.8 |
| REM Onset (min) | 29.3 ± 8.3 | 23.9 ± 4.2 | 35.3 ± 6.0 |
| Wake to NREM (#) | 102.8 ± 6.3 | 107.5 ± 5.9 | **124.8 ± 6.8 \*** |
| NREM to REM (#) | 34.7 ± 2.4 | 30.3 ± 2.1 | 32.3 ± 3.0 |
| REM to Wake (#) | 32.9 ± 2.1 | 29.1 ± 1.9 | 31.6 ± 3.1 |
| NREM to Wake (#) | 70.1 ± 4.6 | 78.3 ± 5.5 | **92.8 ± 8.9 \*** |
| REM to NREM (#) | 1.8 ± 0.4 | 1.2 ± 0.4 | **0.6 ± 0.2 \*** |
| Total Transitions (#) | 242.3 ± 15.3 | 246.4 ± 12.4 | 282.1 ± 12.4 |

| Light Phase | Male |  |  |
| --- | --- | --- | --- |
|  | WT-Control | WT-*Kmo*<sup>+/+</sup> | HET-*Kmo*<sup>+/-</sup> |
| NREM Onset (min) | 6.1 ± 3.0 | 4.6 ± 2.9 | **15.3 ± 2.8 \*** |
| REM Onset (min) | 27.1 ± 3.9 | 20.4 ± 4.1 | 29.2 ± 3.8 |
| Wake to NREM (#) | 112.9 ± 8.1 | 99.1 ± 2.9 | 108.4 ± 6.8 |
| NREM to REM (#) | 34.9 ± 2.2 | 33.3 ± 1.9 | 36.8 ± 2.3 |
| REM to Wake (#) | 32.8 ± 2.1 | 32.1 ± 1.7 | 35.4 ± 2.4 |
| NREM to Wake (#) | 80.1 ± 7.8 | 67.3 ± 3.8 | 72.8 ± 7.6 |
| REM to NREM (#) | 2.1 ± 0.3 | 1.2 ± 0.4 | 1.4 ± 0.4 |
| Total Transitions (#) | 262.9 ± 16.8 | 232.9 ± 5.3 | 254.9 ± 12.9 |

| Dark Phase | Female |  |  |
| --- | --- | --- | --- |
|  | WT-Control | WT-*Kmo*<sup>+/+</sup> | HET-*Kmo*<sup>+/-</sup> |
| NREM Onset (min) | 13.0 ± 4.3 | 13.1 ± 4.4 | 12.0 ± 6.9 |
| REM Onset (min) | 44.9 ± 6.0 | 49.4 ± 13.9 | 59.2 ± 12.5 |
| Wake to NREM (#) | 66.5 ± 5.1 | 68.3 ± 5.7 | 82.1 ± 11.6 |
| NREM to REM (#) | 14.3 ± 1.0 | 16.5 ± 1.0 | 14.8 ± 1.5 |
| REM to Wake (#) | 14.0 ± 0.9 | 16.1 ± 0.9 | 14.3 ± 1.5 |
| NREM to Wake (#) | 52.5 ± 4.9 | 52.2 ± 5.0 | 68.0 ± 11.4 |
| REM to NREM (#) | 0.3 ± 0.2 | 0.3 ± 0.1 | 0.5 ± 0.2 |
| Total Transitions (#) | 147.7 ± 10.3 | 153.4 ± 12.3 | 179.6 ± 23.5 |

| Dark Phase | Male |  |  |
| --- | --- | --- | --- |
|  | WT-Control | WT-*Kmo*<sup>+/+</sup> | HET-*Kmo*<sup>+/-</sup> |
| NREM Onset (min) | 11.7 ± 4.2 | 8.1 ± 2.6 | 13.1 ± 6.6 |
| REM Onset (min) | 101.3 ± 23.4 | 92.2 ± 14.1 | 77.5 ± 14.6 |
| Wake to NREM (#) | 86.5 ± 5.9 | 75.3 ± 4.7 | 77.6 ± 6.0 |
| NREM to REM (#) | 18.1 ± 1.0 | 18.3 ± 1.0 | 19.7 ± 2.0 |
| REM to Wake (#) | 17.6 ± 0.9 | 18.1 ± 0.9 | 19.0 ± 1.9 |
| NREM to Wake (#) | 69.0 ± 5.8 | 57.1 ± 4.5 | 58.7 ± 6.1 |
| REM to NREM (#) | 0.5 ± 0.2 | 0.2 ± 0.1 | 0.7 ± 0.2 |
| Total Transitions (#) | 191.8 ± 12.0 | 168.9 ± 9.8 | 175.6 ± 12.3 |

**Supplementary Figure 1.**
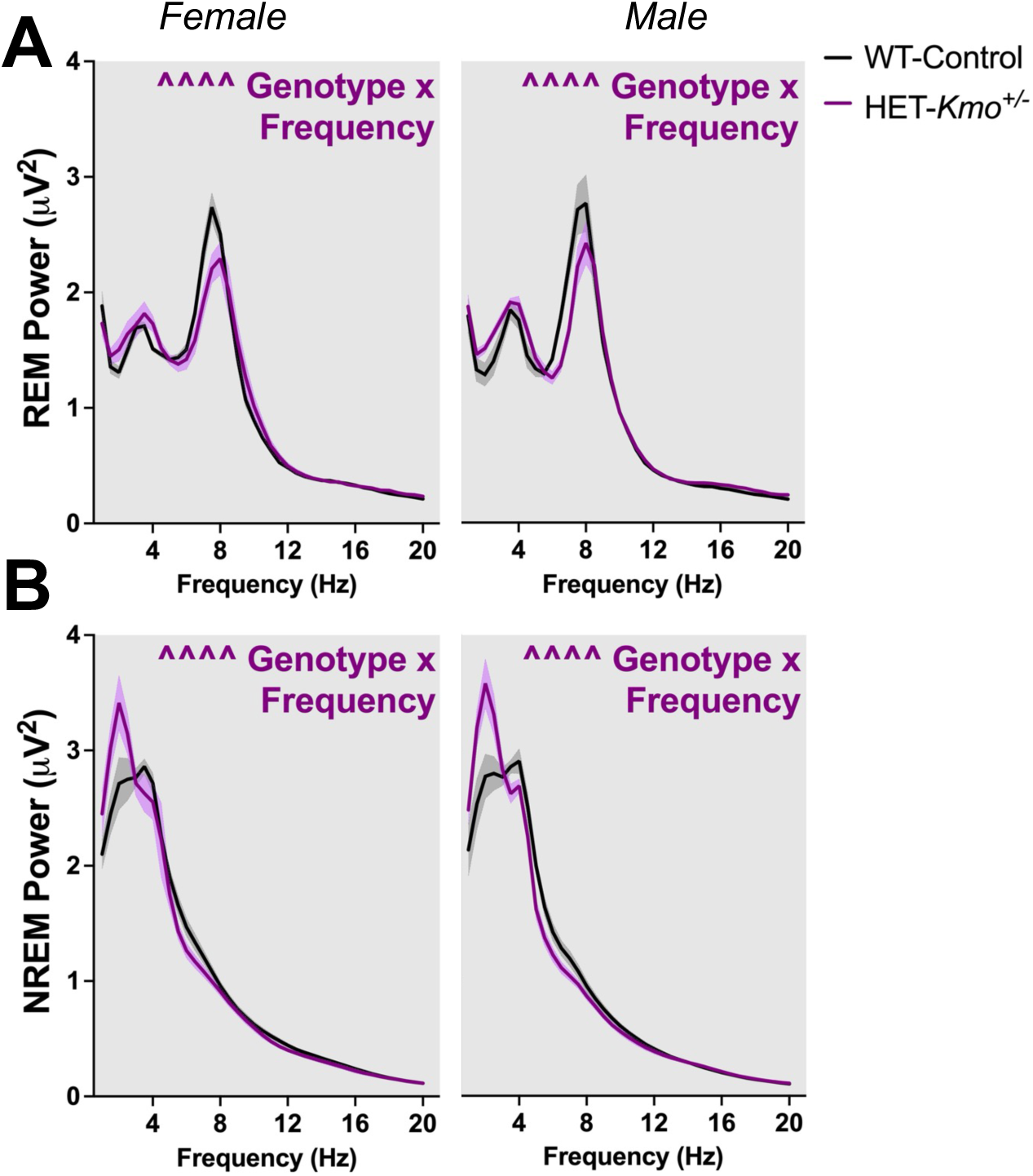
HET-*Kmo^+/−^* mice have altered REM and NREM sleep power spectra in comparison to wild-type (WT-Control) mice during the dark phase. **(A)** REM sleep spectral power (Females: Two-way RM ANOVA Genotype x Frequency interaction F_(38, 380)_= 2.277, ^^^^P<0.0001; Males: Two-way RM ANOVA Genotype x Frequency interaction F_(38, 532)_= 2.765, ^^^^P<0.0001). **(B)** NREM sleep spectral power (Females: Two-way RM ANOVA Genotype x Frequency interaction F_(38, 380)_= 2.455, ^^^^P<0.0001; Males: Two-way RM ANOVA Genotype x Frequency interaction F_(38, 532)_= 4.874, ^^^^P<0.0001). Data are mean ± SEM. N = 6-8 per group.

**Supplementary Figure 2.**
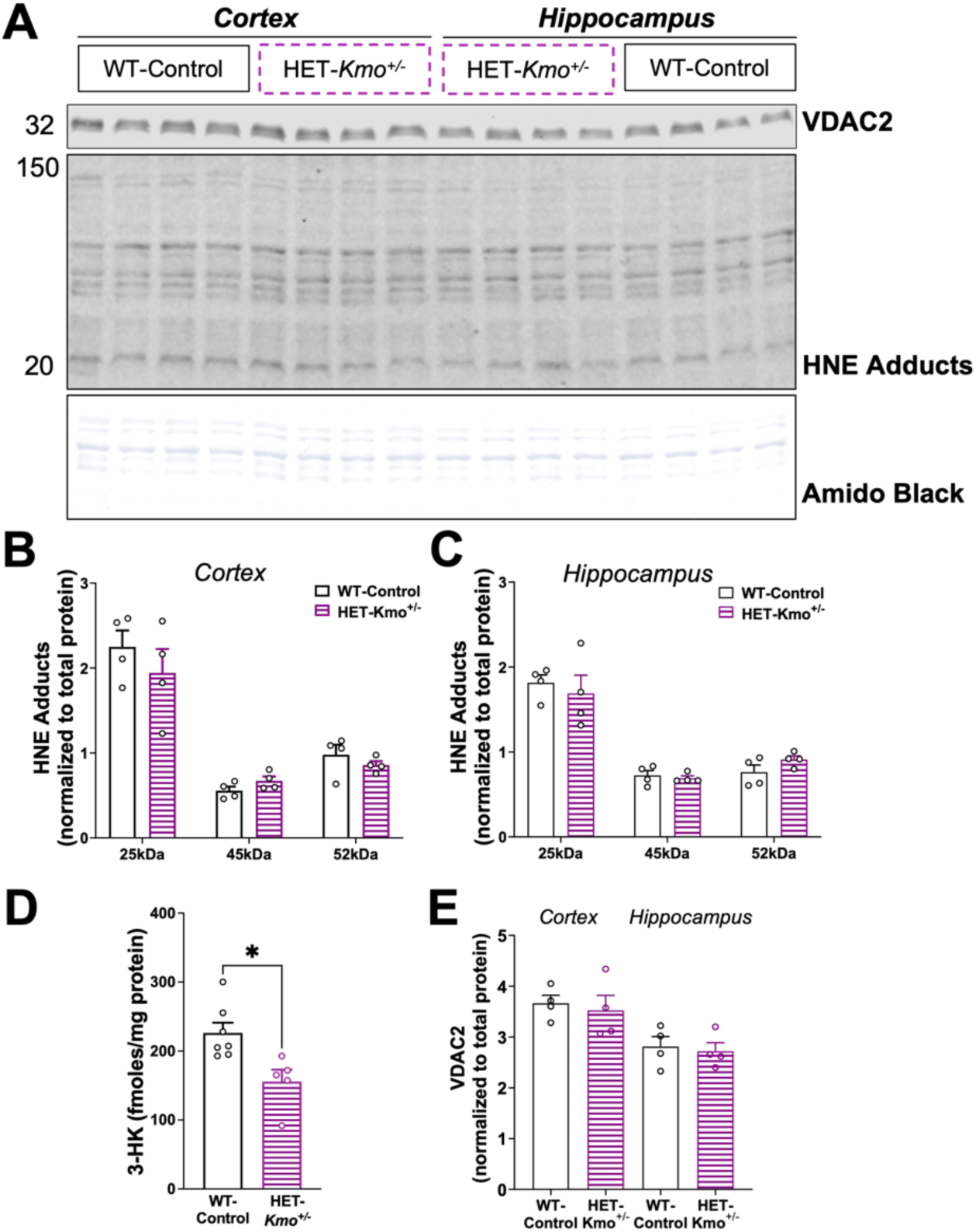
Brain content of 3-hydroxykynurenine is decreased in HET-*Kmo^+/−^* compared to WT-Control female mice. **(A)** Western blot profiling of VDAC2, an outer mitochondrial membrane marker, and HNE adducts, product of lipid peroxidation, in both the cortex and hippocampus of HET-*Kmo^+/−^* female mice and WT-Control. **(B)** HNE adducts in the cortex. **(C)** HNE adducts in the hippocampus. **(D)** 3-hydroxykynurenine (3-HK) in the brain homogenates. **(E)** VDAC2 in the cortical and hippocampal mitochondria. Data are mean ± SEM. Unpaired t test: **P<0.01. N = 4-7 per group.

**Supplementary Figure 3.**
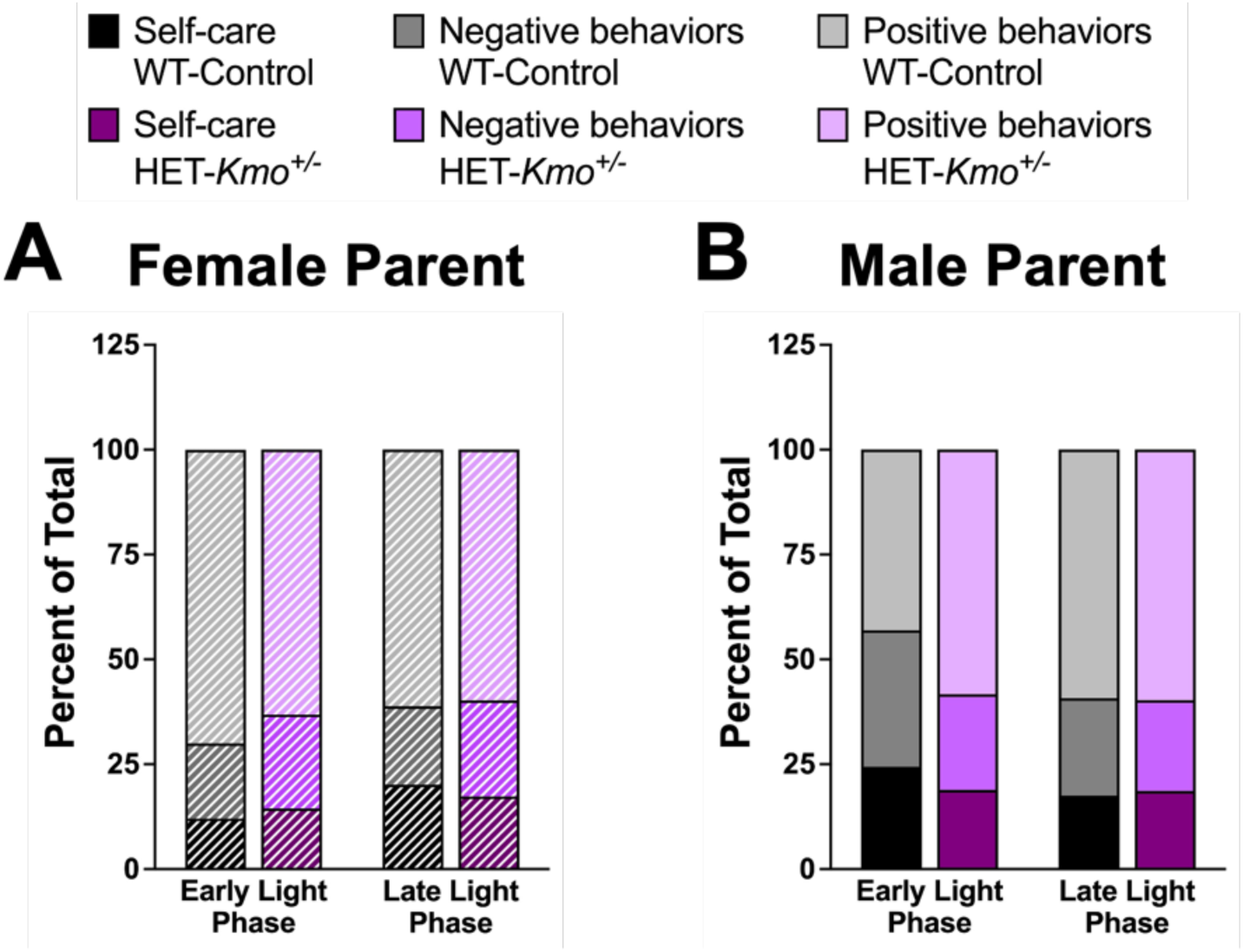
Parental behavior in HET-*Kmo^+/−^* mice in comparison to wild-type (WT-Control) mice. **(A)** Female parent behavior during early and late light phase. **(B)** Male parent behavior during early and late light phase. N = 7-10 per group.

**Supplementary Table 3.** Maternal and paternal behaviors in HET-*Kmo^+/−^* mice in comparison to wild-type (WT-Control) mice during early and late light phase. Data are mean ± SEM. N = 7-10 per group.

| Female Parent Behaviors |  | Positive Behaviors |  |  |
| --- | --- | --- | --- | --- |
|  | # of observations | Grooming pups | Nursing | Contact with pups |
| Early Light Phase | WT-Control | 1.3 ± 0.2 | 7.0 ± 0.5 | 7.8 ± 0.6 |
| | HET-*Kmo*<sup>+/-</sup> | 1.5 ± 0.2 | 6.6 ± 0.8 | 7.5 ± 0.5 |
| Late Light Phase | WT-Control | 1.3 ± 0.1 | 6.5 ± 0.2 | 7.7 ± 0.4 |
| | HET-*Kmo*<sup>+/-</sup> | 1.8 ± 0.3 | 5.0 ± 0.5 | 6.5 ± 0.4 |

| Female Parent Behaviors |  | Self-care |  |
| --- | --- | --- | --- |
|  | # of observations | Self-grooming | Eating and drinking |
| Early Light Phase | WT-Control | 1.5 ± 0.3 | 2.4 ± 0.2 |
| | HET-*Kmo*<sup>+/-</sup> | 1.5 ± 0.5 | 3.3 ± 0.4 |
| Late Light Phase | WT-Control | 1.3 ± 0.2 | 4.2 ± 0.7 |
| | HET-*Kmo*<sup>+/-</sup> | 1.3 ± 0.2 | 3.0 ± 0.3 |

| Female Parent Behaviors |  | Negative Behaviors |  |
| --- | --- | --- | --- |
|  | # of observations | Nest maintenance | Parent out of nest |
| Early Light Phase | WT-Control | 1.7 ± 0.3 | 3.3 ± 1.0 |
| | HET-*Kmo*<sup>+/-</sup> | 1.8 ± 0.8 | 4.4 ± 1.1 |
| Late Light Phase | WT-Control | 1.0 ± 0 | 4.3 ± 0.8 |
| | HET-*Kmo*<sup>+/-</sup> | 1.7 ± 0.2 | 4.1 ± 0.5 |

| Male Parent Behaviors |  | Positive Behaviors |  |
| --- | --- | --- | --- |
|  | # of observations | Grooming pups | Contact with pups |
| Early Light Phase | WT-Control | 1.0 ± 0 | 7.1 ± 0.5 |
| | HET-*Kmo*<sup>+/-</sup> | 1.5 ± 0.5 | 8.0 ± 0.7 |
| Late Light Phase | WT-Control | 1.8 ± 0.3 | 8.9 ± 0.4 |
| | HET-*Kmo*<sup>+/-</sup> | 1.1 ± 0.1 | 8.6 ± 0.2 |

| **Male Parent Behaviors** |  | **Self-care** |  |
| --- | --- | --- | --- |
|  | # of observations | *Self-grooming* | *Eating and drinking* |
| *Early Light Phase* | **WT-Control** | 1.5 ± 0.2 | 3.0 ± 0.3 |
|  | **HET-*Kmo*<sup>+/-</sup>** | 1.5 ± 0.3 | 2.4 ± 0.5 |
| *Late Light Phase* | **WT-Control** | 1.5 ± 0.2 | 2.3 ± 0.5 |
|  | **HET-*Kmo*<sup>+/-</sup>** | 1.8 ± 0.2 | 2.0 ± 0.3 |

| **Male Parent Behaviors** |  | **Negative Behaviors** |  |
| --- | --- | --- | --- |
|  | # of observations | *Nest maintenance* | *Parent out of nest* |
| *Early Light Phase* | **WT-Control** | 1.8 ± 0.4 | 4.3 ± 0.6 |
|  | **HET-*Kmo*<sup>+/-</sup>** | 1.7 ± 0.7 | 3.7 ± 0.6 |
| *Late Light Phase* | **WT-Control** | 1.6 ± 0.5 | 4.1 ± 1.0 |
|  | **HET-*Kmo*<sup>+/-</sup>** | 1.4 ± 0.3 | 2.5 ± 0.2 |

**Supplementary Figure 4.**
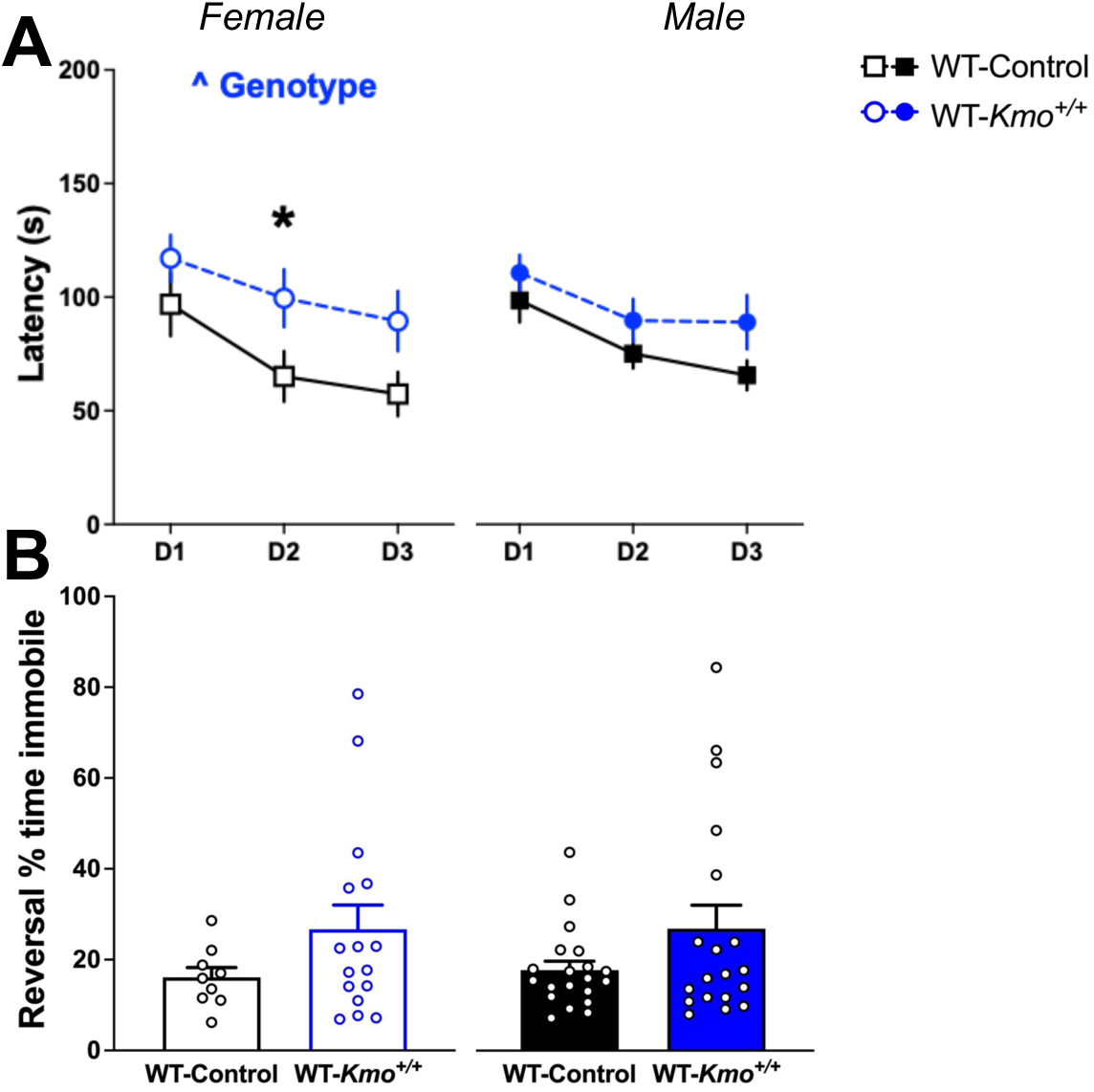
Impaired learning in female wild-type (WT-*Kmo^+/+^*) offspring from HET-*Kmo^+/−^* parents. **(A)** Latency in the Barnes maze (Females: Two-way RM ANOVA Genotype effect F_(1, 36)_= 5.004, ^P<0.05 with Bonferroni’s post hoc test *P<0.05). **(B)** Percent of time immobile in the reversal Barnes maze. Data are mean ± SEM. N = 19-21 per group.

**Supplementary Table 4.** Sleep-wake parameters in WT-*Kmo^+/+^* mice in comparison to wild-type (WT-Control) mice during light and dark phase. Data are mean ± SEM. Unpaired t test: *P<0.05 vs. WT-Control. N = 6-10 per group.

| Light Phase | Female |  | Male |  |
| --- | --- | --- | --- | --- |
|  | WT-Control | WT-*Kmo*<sup>+/+</sup> | WT-Control | WT-*Kmo*<sup>+/+</sup> |
| Total Sleep Duration (min) | 448 ± 9 | **408 ± 14 \*** | 446 ± 10 | 434 ± 8 |
| REM Duration (min) | 40 ± 3 | 35 ± 3 | 41 ± 2 | 43 ± 2 |
| REM Bouts (#) | 35 ± 3 | 31 ± 2 | 36 ± 2 | 34 ± 2 |
| NREM Bouts (#) | 111 ± 7 | 114 ± 6 | 121 ± 8 | 107 ± 3 |
| Wake Bouts (#) | 107 ± 6 | 113 ± 6 | 118 ± 9 | 104 ± 3 |
| REM Bout Duration (s/bout) | 71 ± 2 | 69 ± 2 | 75 ± 2 | 81 ± 2 |
| NREM Bout Duration (s/bout) | 235 ± 19 | 208 ± 7 | 227 ± 29 | 237 ± 10 |
| Wake Bout Duration (s/bout) | 196 ± 14 | 235 ± 35 | 189 ± 16 | **238 ± 15 \*** |

| Dark Phase | Female |  | Male |  |
| --- | --- | --- | --- | --- |
|  | WT-Control | WT-*Kmo*<sup>+/+</sup> | WT-Control | WT-*Kmo*<sup>+/+</sup> |
| Total Sleep Duration (min) | 265 ± 20 | 262 ± 16 | 281 ± 16 | 277 ± 12 |
| REM Duration (min) | 17 ± 1 | 19 ± 2 | 21 ± 1 | 20 ± 1 |
| NREM Duration (min) | 248 ± 20 | 242 ± 16 | 260 ± 15 | 257 ± 11 |
| Wake Duration (min) | 455 ± 20 | 458 ± 16 | 439 ± 16 | 443 ± 12 |
| REM Bouts (#) | 15 ± 1 | 17 ± 1 | 18 ± 1 | 18 ± 1 |
| NREM Bouts (#) | 71 ± 5 | 72 ± 6 | 92 ± 6 | 80 ± 5 |
| Wake Bouts (#) | 74 ± 5 | 76 ± 5 | 93 ± 6 | 82 ± 5 |
| REM Bout Duration (s/bout) | 77 ± 5 | 70 ± 2 | 72 ± 3 | 67 ± 3 |
| NREM Bout Duration (s/bout) | 223 ± 24 | 202 ± 3 | 178 ± 12 | 196 ± 11 |
| Wake Bout Duration (s/bout) | 769 ± 92 | 770 ± 128 | 619 ± 127 | 688 ± 61 |

**Supplementary Figure 5.**
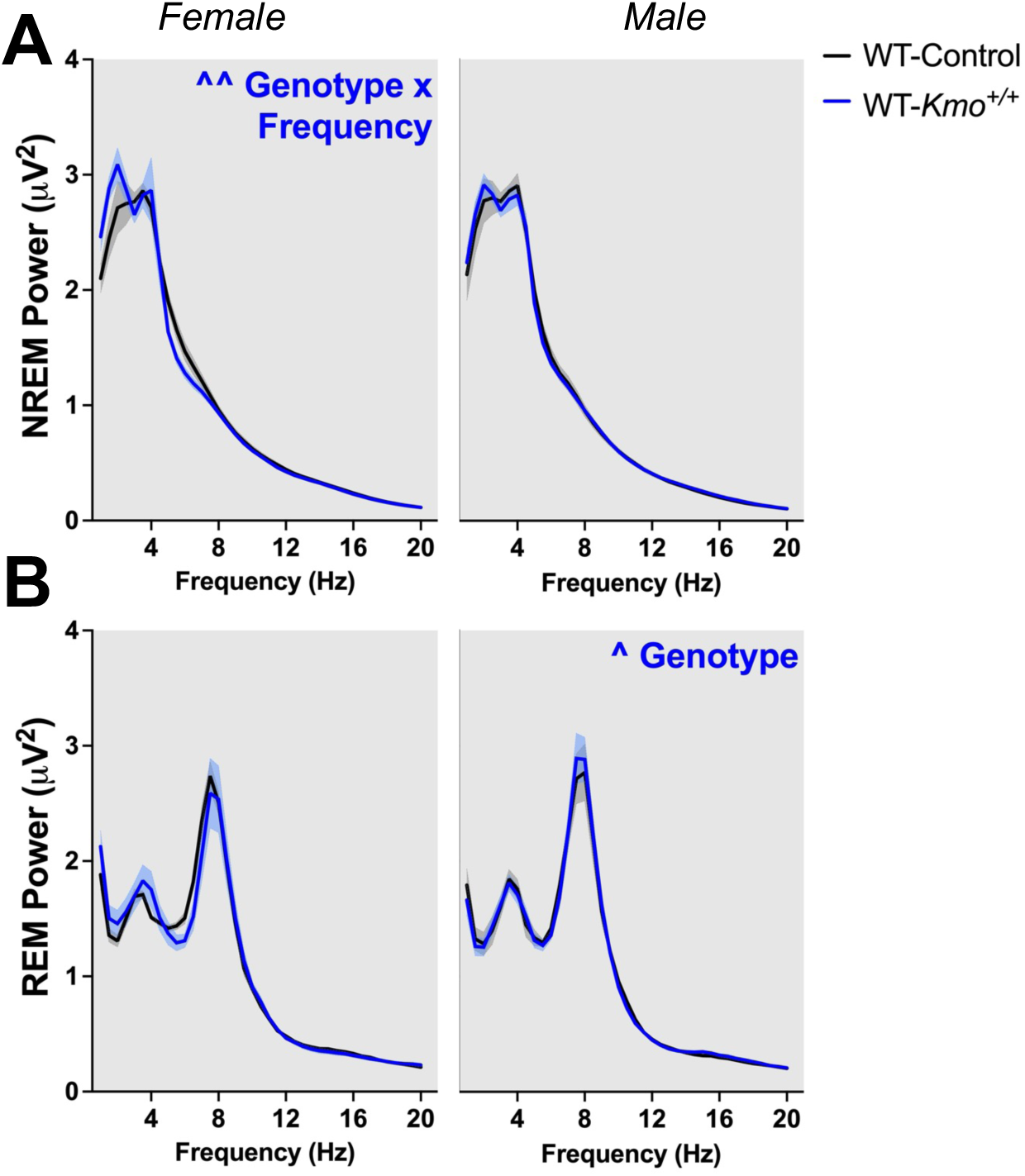
Sex-specific differences in NREM and REM sleep power spectra in WT-*Kmo^+/+^* compared to wild-type (WT-Control) mice during the dark phase. **(A)** NREM sleep spectral power (Females: Two-way RM ANOVA Genotype x Frequency interaction F_(38, 380)_= 1.821, ^^P<0.01). **(B)** REM sleep spectral power (Males: Two-way RM ANOVA Genotype effect F_(1, 16)_= 4.676, ^P<0.05). Data are mean ± SEM. N = 6-10 per group.

## REFERENCES

1. Bai D, Yip BHK, Windham GC, Sourander A, Francis R, Yoffe R, et al. Association of Genetic and Environmental Factors With Autism in a 5-Country Cohort. JAMA Psychiatry. 2019;76:10:1035; doi:10.1001/jamapsychiatry.2019.1411.

2. Brikell I, Kuja-Halkola R, Larsson H. Heritability of attention-deficit hyperactivity disorder in adults. American Journal of Medical Genetics Part B: Neuropsychiatric Genetics. 2015;168:6:406–13; doi:10.1002/ajmg.b.32335.

3. Smoller JW, Finn CT. Family, twin, and adoption studies of bipolar disorder. American Journal of Medical Genetics Part C: Seminars in Medical Genetics. 2003;123C:1:48–58; doi:10.1002/ajmg.c.20013.

4. Sullivan PF, Kendler KS, Neale MC. Schizophrenia as a Complex Trait. Archives of General Psychiatry. 2003;60:12:1187; doi:10.1001/archpsyc.60.12.1187.

5. Genetic relationship between five psychiatric disorders estimated from genome-wide SNPs. Nature Genetics. 2013;45:9:984–94; doi:10.1038/ng.2711.

6. Derks EM, Thorp JG, Gerring ZF. Ten challenges for clinical translation in psychiatric genetics. Nature Genetics. 2022;54:10:1457–65; doi:10.1038/s41588-022-01174-0.

7. Lee PH, Feng Y-CA, Smoller JW. Pleiotropy and Cross-Disorder Genetics Among Psychiatric Disorders. Biological Psychiatry. 2021;89:1:20–31; doi:10.1016/j.biopsych.2020.09.026.

8. Bölte S, Neufeld J, Marschik PB, Williams ZJ, Gallagher L, Lai M-C. Sex and gender in neurodevelopmental conditions. Nature Reviews Neurology. 2023;19:3:136–59; doi:10.1038/s41582-023-00774-6.

9. May T, Adesina I, McGillivray J, Rinehart NJ. Sex differences in neurodevelopmental disorders. Current Opinion in Neurology. 2019;32:4:622–6; doi:10.1097/wco.0000000000000714.

10. Surén P, Bakken IJ, Aase H, Chin R, Gunnes N, Lie KK, et al. Autism Spectrum Disorder, ADHD, Epilepsy, and Cerebral Palsy in Norwegian Children. Pediatrics. 2012;130:1:e152–e8; doi:10.1542/peds.2011-3217.

11. Sommer IE, Tiihonen J, van Mourik A, Tanskanen A, Taipale H. The clinical course of schizophrenia in women and men—a nation-wide cohort study. npj Schizophrenia. 2020;6:1; doi:10.1038/s41537-020-0102-z.

12. Erhardt S, Pocivavsek A, Repici M, Liu X-C, Imbeault S, Maddison DC, et al. Adaptive and Behavioral Changes in Kynurenine 3-Monooxygenase Knockout Mice: Relevance to Psychotic Disorders. Biological Psychiatry. 2017;82:10:756–65; doi:10.1016/j.biopsych.2016.12.011.

13. Lim CK, Essa MM, de Paula Martins R, Lovejoy DB, Bilgin AA, Waly MI, et al. Altered kynurenine pathway metabolism in autism: Implication for immune-induced glutamatergic activity. Autism Research. 2015;9:6:621–31; doi:10.1002/aur.1565.

14. Linderholm KR, Skogh E, Olsson SK, Dahl ML, Holtze M, Engberg G, et al. Increased Levels of Kynurenine and Kynurenic Acid in the CSF of Patients With Schizophrenia. Schizophrenia Bulletin. 2010;38:3:426–32; doi:10.1093/schbul/sbq086.

15. Oades RD, Dauvermann MR, Schimmelmann BG, Schwarz MJ, Myint A-M. Attention-deficit hyperactivity disorder (ADHD) and glial integrity: S100B, cytokines and kynurenine metabolism - effects of medication. Behavioral and Brain Functions. 2010;6:1:29; doi:10.1186/1744-9081-6-29.

16. Olsson SK, Samuelsson M, Saetre P, Lindström L, Jönsson EG, Nordin C, et al. Elevated levels of kynurenic acid in the cerebrospinal fluid of patients with bipolar disorder. Journal of Psychiatry and Neuroscience. 2010;35:3:195–9; doi:10.1503/jpn.090180.

17. Schwarcz R, Rassoulpour A, Wu HQ, Medoff D, Tamminga CA, Roberts RC. Increased cortical kynurenate content in schizophrenia. Biol Psychiatry. 2001;50:7:521–30; doi:10.1016/s0006-3223(01)01078-2.

18. Kegel ME, Johansson V, Wetterberg L, Bhat M, Schwieler L, Cannon TD, et al. Kynurenic acid and psychotic symptoms and personality traits in twins with psychiatric morbidity. Psychiatry Res. 2017;247:105–12; doi:10.1016/j.psychres.2016.11.017.

19. Antenucci N, D’Errico G, Fazio F, Nicoletti F, Bruno V, Battaglia G. Changes in kynurenine metabolites in the gray and white matter of the dorsolateral prefrontal cortex of individuals affected by schizophrenia. Schizophrenia (Heidelb). 2024;10:1:27; doi:10.1038/s41537-024-00447-3.

20. Comai S, Bertazzo A, Vachon J, Daigle M, Toupin J, Cote G, et al. Tryptophan via serotonin/kynurenine pathways abnormalities in a large cohort of aggressive inmates: markers for aggression. Prog Neuropsychopharmacol Biol Psychiatry. 2016;70:8–16; doi:10.1016/j.pnpbp.2016.04.012.

21. Beggiato S, Notarangelo FM, Sathyasaikumar KV, Giorgini F, Schwarcz R. Maternal genotype determines kynurenic acid levels in the fetal brain: Implications for the pathophysiology of schizophrenia. Journal of Psychopharmacology. 2018;32:11:1223–32; doi:10.1177/0269881118805492.

22. McFarlane HG, Kusek GK, Yang M, Phoenix JL, Bolivar VJ, Crawley JN. Autism-like behavioral phenotypes in BTBR T+tf/J mice. Genes, Brain and Behavior. 2007;7:2:152–63; doi:10.1111/j.1601-183x.2007.00330.x.

23. Murakami Y, Imamura Y, Saito K, Sakai D, Motoyama J. Altered kynurenine pathway metabolites in a mouse model of human attention-deficit hyperactivity/autism spectrum disorders: A potential new biological diagnostic marker. Scientific Reports. 2019;9:1; doi:10.1038/s41598-019-49781-y.

24. Notarangelo FM, Pocivavsek A. Elevated kynurenine pathway metabolism during neurodevelopment: Implications for brain and behavior. Neuropharmacology. 2017;112:275–85; doi:10.1016/j.neuropharm.2016.03.001.

25. Cervenka I, Agudelo LZ, Ruas JL. Kynurenines: Tryptophan’s metabolites in exercise, inflammation, and mental health. Science. 2017;357:6349; doi:10.1126/science.aaf9794.

26. Wonodi I, McMahon RP, Krishna N, Mitchell BD, Liu J, Glassman M, et al. Influence of kynurenine 3-monooxygenase (KMO) gene polymorphism on cognitive function in schizophrenia. Schizophrenia Research. 2014;160:1-3:80–7; doi:10.1016/j.schres.2014.10.026.

27. Wonodi I, Stine OC, Sathyasaikumar KV, Roberts RC, Mitchell BD, Hong LE, et al. Downregulated Kynurenine 3-Monooxygenase Gene Expression and Enzyme Activity in Schizophrenia and Genetic Association With Schizophrenia Endophenotypes. Archives of General Psychiatry. 2011;68:7:665; doi:10.1001/archgenpsychiatry.2011.71.

28. Sathyasaikumar KV, Stachowski EK, Wonodi I, Roberts RC, Rassoulpour A, McMahon RP, et al. Impaired Kynurenine Pathway Metabolism in The Prefrontal Cortex of Individuals With Schizophrenia. Schizophrenia Bulletin. 2010;37:6:1147–56; doi:10.1093/schbul/sbq112.

29. Holtze M, Saetre P, Engberg G, Schwieler L, Werge T, Andreassen OA, et al. Kynurenine 3-monooxygenase polymorphisms: relevance for kynurenic acid synthesis in patients with schizophrenia and healthy controls. Journal of Psychiatry and Neuroscience. 2012;37:1:53–7; doi:10.1503/jpn.100175.

30. Lavebratt C, Olsson S, Backlund L, Frisén L, Sellgren C, Priebe L, et al. The KMO allele encoding Arg452 is associated with psychotic features in bipolar disorder type 1, and with increased CSF KYNA level and reduced KMO expression. Molecular Psychiatry. 2013;19:3:334–41; doi:10.1038/mp.2013.11.

31. Bortz DM, Wu HQ, Schwarcz R, Bruno JP. Oral administration of a specific kynurenic acid synthesis (KAT II) inhibitor attenuates evoked glutamate release in rat prefrontal cortex. Neuropharmacology. 2017;121:69–78; doi:10.1016/j.neuropharm.2017.04.023.

32. Hilmas C, Pereira EFR, Alkondon M, Rassoulpour A, Schwarcz R, Albuquerque EX. The Brain Metabolite Kynurenic Acid Inhibits α7 Nicotinic Receptor Activity and Increases Non-α7 Nicotinic Receptor Expression: Physiopathological Implications. The Journal of Neuroscience. 2001;21:19:7463–73; doi:10.1523/jneurosci.21-19-07463.2001.

33. Pocivavsek A, Baratta AM, Mong JA, Viechweg SS. Acute Kynurenine Challenge Disrupts Sleep–Wake Architecture and Impairs Contextual Memory in Adult Rats. Sleep. 2017;40:11; doi:10.1093/sleep/zsx141.

34. Alexander KS, Wu HQ, Schwarcz R, Bruno JP. Acute elevations of brain kynurenic acid impair cognitive flexibility: normalization by the alpha7 positive modulator galantamine. Psychopharmacology (Berl). 2012;220:3:627–37; doi:10.1007/s00213-011-2539-2.

35. Pocivavsek A, Wu HQ, Potter MC, Elmer GI, Pellicciari R, Schwarcz R. Fluctuations in endogenous kynurenic acid control hippocampal glutamate and memory. Neuropsychopharmacology. 2011;36:11:2357–67; doi:10.1038/npp.2011.127.

36. Pocivavsek A, Notarangelo FM, Wu HQ, Bruno JP, Schwarcz R. Astrocytes as pharmacological targets in the treatment of schizophrenia: Focus on kynurenic acid. In: Pletnikov MV, Waddinton J, editors. Modeling the psychopathological dimensions of schizophrenia: From molecules to behavior Elsevier Academic Press; 2016. p. 423–43.

37. Wu HQ, Pereira EF, Bruno JP, Pellicciari R, Albuquerque EX, Schwarcz R. The astrocyte-derived alpha7 nicotinic receptor antagonist kynurenic acid controls extracellular glutamate levels in the prefrontal cortex. J Mol Neurosci. 2010;40:1-2:204–10; doi:10.1007/s12031-009-9235-2.

38. Parenti I, Rabaneda LG, Schoen H, Novarino G. Neurodevelopmental Disorders: From Genetics to Functional Pathways. Trends in Neurosciences. 2020;43:8:608–21; doi:10.1016/j.tins.2020.05.004.

39. Shelton AR, Malow B. Neurodevelopmental Disorders Commonly Presenting with Sleep Disturbances. Neurotherapeutics. 2021;18:1:156–69; doi:10.1007/s13311-020-00982-8.

40. Pocivavsek A, Erhardt S. Kynurenic acid: translational perspectives of a therapeutically targetable gliotransmitter. Neuropsychopharmacology. 2024;49:1:307–8; doi:10.1038/s41386-023-01681-6.

41. Forrest CM, Khalil OS, Pisar M, Darlington LG, Stone TW. Prenatal inhibition of the tryptophan-kynurenine pathway alters synaptic plasticity and protein expression in the rat hippocampus. Brain research. 2013;1504:1–15; doi:10.1016/j.brainres.2013.01.031.

42. Pocivavsek A, Thomas MAR, Elmer GI, Bruno JP, Schwarcz R. Continuous kynurenine administration during the prenatal period, but not during adolescence, causes learning and memory deficits in adult rats. Psychopharmacology. 2014;231:14:2799–809; doi:10.1007/s00213-014-3452-2.

43. Goeden N, Notarangelo FM, Pocivavsek A, Beggiato S, Bonnin A, Schwarcz R. Prenatal Dynamics of Kynurenine Pathway Metabolism in Mice: Focus on Kynurenic Acid. Developmental Neuroscience. 2017;39:6:519–28; doi:10.1159/000481168.

44. Rentschler KM, Baratta AM, Ditty AL, Wagner NTJ, Wright CJ, Milosavljevic S, et al. Prenatal Kynurenine Elevation Elicits Sex-Dependent Changes in Sleep and Arousal During Adulthood: Implications for Psychotic Disorders. Schizophrenia Bulletin. 2021;47:5:1320–30; doi:10.1093/schbul/sbab029.

45. Buck SA, Baratta AM, Pocivavsek A. Exposure to elevated embryonic kynurenine in rats: Sex-dependent learning and memory impairments in adult offspring. Neurobiology of Learning and Memory. 2020;174:107282; doi:10.1016/j.nlm.2020.107282.

46. Pershing ML, Bortz DM, Pocivavsek A, Fredericks PJ, Jorgensen CV, Vunck SA, et al. Elevated levels of kynurenic acid during gestation produce neurochemical, morphological, and cognitive deficits in adulthood: implications for schizophrenia. Neuropharmacology. 2015;90:33–41; doi:10.1016/j.neuropharm.2014.10.017.

47. Richardson RB, Mailloux RJ. Mitochondria Need Their Sleep: Redox, Bioenergetics, and Temperature Regulation of Circadian Rhythms and the Role of Cysteine-Mediated Redox Signaling, Uncoupling Proteins, and Substrate Cycles. Antioxidants. 2023;12:3:674; doi:10.3390/antiox12030674.

48. Tanaka D, Nakada K, Takao K, Ogasawara E, Kasahara A, Sato A, et al. Normal mitochondrial respiratory function is essential for spatial remote memory in mice. Molecular Brain. 2008;1:1; doi:10.1186/1756-6606-1-21.

49. Underwood EL, Redell JB, Hood KN, Maynard ME, Hylin M, Waxham MN, et al. Enhanced presynaptic mitochondrial energy production is required for memory formation. Scientific Reports. 2023;13:1; doi:10.1038/s41598-023-40877-0.

50. Hirai K, Kuroyanagi H, Tatebayashi Y, Hayashi Y, Hirabayashi-Takahashi K, Saito K, et al. Dual role of the carboxyl-terminal region of pig liver L-kynurenine 3-monooxygenase: mitochondrial-targeting signal and enzymatic activity. Journal of Biochemistry. 2010;148:6:639–50; doi:10.1093/jb/mvq099.

51. Maddison DC, Alfonso-Núñez M, Swaih AM, Breda C, Campesan S, Allcock N, et al. A novel role for kynurenine 3-monooxygenase in mitochondrial dynamics. PLOS Genetics. 2020;16:11:e1009129; doi:10.1371/journal.pgen.1009129.

52. Sadakata M, Fujii K, Kaneko R, Hosoya E, Sugimoto H, Kawabata-Iwakawa R, et al. Maternal immunoglobulin G affects brain development of mouse offspring. Journal of Neuroinflammation. 2024;21:1; doi:10.1186/s12974-024-03100-z.

53. Giorgini F, Huang S-Y, Sathyasaikumar KV, Notarangelo FM, Thomas MAR, Tararina M, et al. Targeted Deletion of Kynurenine 3-Monooxygenase in Mice. Journal of Biological Chemistry. 2013;288:51:36554–66; doi:10.1074/jbc.m113.503813.

54. Sambrook J, Russell DW. Purification of RNA from Cells and Tissues by Acid Phenol-Guanidinium Thiocyanate-Chloroform Extraction. Cold Spring Harbor Protocols. 2006;2006:1:pdb.prot4045; doi:10.1101/pdb.prot4045.

55. Muranishi Y, Parry L, Averous J, Terrisse A, Maurin A-C, Chaveroux C, et al. Method for Collecting Mouse Milk without Exogenous Oxytocin Stimulation. BioTechniques. 2016;60:1:47–9; doi:10.2144/000114373.

56. Notarangelo Francesca M, Beggiato S, Schwarcz R. Assessment of Prenatal Kynurenine Metabolism Using Tissue Slices: Focus on the Neosynthesis of Kynurenic Acid in Mice. Developmental Neuroscience. 2019;41:1-2:102–11; doi:10.1159/000499736.

57. Sbornova I, van der Sande E, Milosavljevic S, Amurrio E, Burbano SD, Das PK, et al. The Sleep Quality- and Myopia-Linked PDE11A-Y727C Variant Impacts Neural Physiology by Reducing Catalytic Activity and Altering Subcellular Compartmentalization of the Enzyme. Cells. 2023;12:24:2839; doi:10.3390/cells12242839.

58. Sperling JA, Sakamuri SSVP, Albuck AL, Sure VN, Evans WR, Peterson NR, et al. Measuring Respiration in Isolated Murine Brain Mitochondria: Implications for Mechanistic Stroke Studies. NeuroMolecular Medicine. 2019;21:4:493–504; doi:10.1007/s12017-019-08552-8.

59. Lowry O, Rosebrough N, Farr AL, Randall R. PROTEIN MEASUREMENT WITH THE FOLIN PHENOL REAGENT. Journal of Biological Chemistry. 1951;193:1:265–75; doi:10.1016/s0021-9258(19)52451-6.

60. Barnes CA. Memory deficits associated with senescence: A neurophysiological and behavioral study in the rat. Journal of Comparative and Physiological Psychology. 1979;93:1:74–104; doi:10.1037/h0077579.

61. Tucker LB, McCabe JT. Behavior of Male and Female C57BL/6J Mice Is More Consistent with Repeated Trials in the Elevated Zero Maze than in the Elevated Plus Maze. Frontiers in Behavioral Neuroscience. 2017;11; doi:10.3389/fnbeh.2017.00013.

62. Calpe-López C, García-Pardo MP, Martínez-Caballero MA, Santos-Ortíz A, Aguilar MA. Behavioral Traits Associated With Resilience to the Effects of Repeated Social Defeat on Cocaine-Induced Conditioned Place Preference in Mice. Frontiers in Behavioral Neuroscience. 2020;13; doi:10.3389/fnbeh.2019.00278.

63. Quadir SG, Arleth GM, Jahad JV, Echeveste Sanchez M, Effinger DP, Herman MA. Sex differences in affective states and association with voluntary ethanol intake in Sprague–Dawley rats. Psychopharmacology. 2022;239:2:589–604; doi:10.1007/s00213-021-06052-x.

64. Chourbaji S, Hoyer C, Richter SH, Brandwein C, Pfeiffer N, Vogt MA, et al. Differences in Mouse Maternal Care Behavior – Is There a Genetic Impact of the Glucocorticoid Receptor? PLoS ONE. 2011;6:4:e19218; doi:10.1371/journal.pone.0019218.

65. Orso R, Wearick-Silva LE, Creutzberg KC, Centeno-Silva A, Glusman Roithmann L, Pazzin R, et al. Maternal behavior of the mouse dam toward pups: implications for maternal separation model of early life stress. Stress. 2017;21:1:19–27; doi:10.1080/10253890.2017.1389883.

66. Rentschler KM, Milosavljevic S, Baratta AM, Wright CJ, Piroli MV, Tentor Z, et al. Reducing brain kynurenic acid synthesis precludes kynurenine-induced sleep disturbances. J Sleep Res. 2024;33:3:e14038; doi:10.1111/jsr.14038.

67. Milosavljevic S, Smith AK, Wright CJ, Valafar H, Pocivavsek A. Kynurenine aminotransferase II inhibition promotes sleep and rescues impairments induced by neurodevelopmental insult. Transl Psychiatry. 2023;13:1:106; doi:10.1038/s41398-023-02399-1.

68. Hartmann C, Kempf A. Mitochondrial control of sleep. Current Opinion in Neurobiology. 2023;81:102733; doi:10.1016/j.conb.2023.102733.

69. Xian H, Liou Y-C. Functions of outer mitochondrial membrane proteins: mediating the crosstalk between mitochondrial dynamics and mitophagy. Cell Death &amp; Differentiation. 2020;28:3:827–42; doi:10.1038/s41418-020-00657-z.

70. Bhatti JS, Bhatti GK, Reddy PH. Mitochondrial dysfunction and oxidative stress in metabolic disorders — A step towards mitochondria based therapeutic strategies. Biochimica et Biophysica Acta (BBA) - Molecular Basis of Disease. 2017;1863:5:1066–77; doi:10.1016/j.bbadis.2016.11.010.

71. Guo C, Sun L, Chen X, Zhang D. Oxidative stress, mitochondrial damage and neurodegenerative diseases. Neural Regen Res. 2013;8:21:2003–14; doi:10.3969/j.issn.1673-5374.2013.21.009.

72. Haynes PR, Pyfrom ES, Li Y, Stein C, Cuddapah VA, Jacobs JA, et al. A neuron–glia lipid metabolic cycle couples daily sleep to mitochondrial homeostasis. Nature Neuroscience. 2024;27:4:666–78; doi:10.1038/s41593-023-01568-1.

73. Singh A, Kukreti R, Saso L, Kukreti S. Oxidative Stress: A Key Modulator in Neurodegenerative Diseases. Molecules. 2019;24:8:1583; doi:10.3390/molecules24081583.

74. Dalleau S, Baradat M, Guéraud F, Huc L. Cell death and diseases related to oxidative stress:4-hydroxynonenal (HNE) in the balance. Cell Death &amp; Differentiation. 2013;20:12:1615–30; doi:10.1038/cdd.2013.138.

75. Okuda S, Nishiyama N, Saito H, Katsuki H. 3-Hydroxykynurenine, an Endogenous Oxidative Stress Generator, Causes Neuronal Cell Death with Apoptotic Features and Region Selectivity. Journal of Neurochemistry. 1998;70:1:299–307; doi:10.1046/j.1471-4159.1998.70010299.x.

76. Sas K, Szabó E, Vécsei L. Mitochondria, Oxidative Stress and the Kynurenine System, with a Focus on Ageing and Neuroprotection. Molecules. 2018;23:1:191; doi:10.3390/molecules23010191.

77. Gawel K, Gibula E, Marszalek-Grabska M, Filarowska J, Kotlinska JH. Assessment of spatial learning and memory in the Barnes maze task in rodents—methodological consideration. Naunyn-Schmiedeberg’s Archives of Pharmacology. 2018;392:1:1–18; doi:10.1007/s00210-018-1589-y.

78. Negrón-Oyarzo I, Espinosa N, Aguilar-Rivera M, Fuenzalida M, Aboitiz F, Fuentealba P. Coordinated prefrontal–hippocampal activity and navigation strategy-related prefrontal firing during spatial memory formation. Proceedings of the National Academy of Sciences. 2018;115:27:7123–8; doi:10.1073/pnas.1720117115.

79. Diekelmann S, Born J. The memory function of sleep. Nat Rev Neurosci. 2010;11:2:114–26; doi:10.1038/nrn2762.

80. Graves L, Pack A, Abel T. Sleep and memory: a molecular perspective. Trends Neurosci. 2001;24:4:237–43; doi:10.1016/s0166-2236(00)01744-6.

81. Rasch B, Born J. About Sleep’s Role in Memory. Physiological Reviews. 2013;93:2:681–766; doi:10.1152/physrev.00032.2012.

82. Jami ES, Hammerschlag AR, Bartels M, Middeldorp CM. Parental characteristics and offspring mental health and related outcomes: a systematic review of genetically informative literature. Translational Psychiatry. 2021;11:1; doi:10.1038/s41398-021-01300-2.

83. Kong A, Thorleifsson G, Frigge ML, Vilhjalmsson BJ, Young AI, Thorgeirsson TE, et al. The nature of nurture: Effects of parental genotypes. Science. 2018;359:6374:424–8; doi:10.1126/science.aan6877.

84. Knop J, Joëls M, van der Veen R. The added value of rodent models in studying parental influence on offspring development: opportunities, limitations and future perspectives. Current Opinion in Psychology. 2017;15:174–81; doi:10.1016/j.copsyc.2017.02.030.

85. Brandon NJ, Sawa A. Linking neurodevelopmental and synaptic theories of mental illness through DISC1. Nature Reviews Neuroscience. 2011;12:12:707–22; doi:10.1038/nrn3120.

86. de la Torre-Ubieta L, Won H, Stein JL, Geschwind DH. Advancing the understanding of autism disease mechanisms through genetics. Nature Medicine. 2016;22:4:345–61; doi:10.1038/nm.4071.

87. Pardo CA, Eberhart CG. The Neurobiology of Autism. Brain Pathology. 2007;17:4:434–47; doi:10.1111/j.1750-3639.2007.00102.x.

88. Ross CA, Margolis RL, Reading SAJ, Pletnikov M, Coyle JT. Neurobiology of Schizophrenia. Neuron. 2006;52:1:139–53; doi:10.1016/j.neuron.2006.09.015.

89. Zoghbi HY, Bear MF. Synaptic Dysfunction in Neurodevelopmental Disorders Associated with Autism and Intellectual Disabilities. Cold Spring Harbor Perspectives in Biology. 2012;4:3:a009886-a; doi:10.1101/cshperspect.a009886.

90. Tufvesson-Alm M, Schwieler L, Schwarcz R, Goiny M, Erhardt S, Engberg G. Importance of kynurenine 3-monooxygenase for spontaneous firing and pharmacological responses of midbrain dopamine neurons: Relevance for schizophrenia. Neuropharmacology. 2018;138:130–9; doi:10.1016/j.neuropharm.2018.06.003.

91. Erhardt S, Blennow K, Nordin C, Skogh E, Lindstrom LH, Engberg G. Kynurenic acid levels are elevated in the cerebrospinal fluid of patients with schizophrenia. Neuroscience letters. 2001;313:1-2:96–8; http://www.ncbi.nlm.nih.gov/pubmed/11684348.

92. Linderholm KR, Skogh E, Olsson SK, Dahl ML, Holtze M, Engberg G, et al. Increased levels of kynurenine and kynurenic acid in the CSF of patients with schizophrenia. Schizophr Bull. 2012;38:3:426–32; doi:10.1093/schbul/sbq086.

93. Nilsson LK, Linderholm KR, Engberg G, Paulson L, Blennow K, Lindstrom LH, et al. Elevated levels of kynurenic acid in the cerebrospinal fluid of male patients with schizophrenia. Schizophr Res. 2005;80:2-3:315–22; doi:10.1016/j.schres.2005.07.013.

94. Sellgren CM, Kegel ME, Bergen SE, Ekman CJ, Olsson S, Larsson M, et al. A genome-wide association study of kynurenic acid in cerebrospinal fluid: implications for psychosis and cognitive impairment in bipolar disorder. Mol Psychiatry. 2016;21:10:1342–50; doi:10.1038/mp.2015.186.

95. Chess AC, Simoni MK, Alling TE, Bucci DJ. Elevations of endogenous kynurenic acid produce spatial working memory deficits. Schizophr Bull. 2007;33:3:797–804; doi:10.1093/schbul/sbl033.

96. Erhardt S, Schwieler L, Emanuelsson C, Geyer M. Endogenous kynurenic acid disrupts prepulse inhibition. Biol Psychiatry. 2004;56:4:255–60; doi:10.1016/j.biopsych.2004.06.006.

97. DiNuzzo M, Nedergaard M. Brain energetics during the sleep–wake cycle. Current Opinion in Neurobiology. 2017;47:65–72; doi:10.1016/j.conb.2017.09.010.

98. Aalling NN, Nedergaard M, DiNuzzo M. Cerebral Metabolic Changes During Sleep. Curr Neurol Neurosci Rep. 2018;18:9:57; doi:10.1007/s11910-018-0868-9.

99. Dworak M, McCarley RW, Kim T, Kalinchuk AV, Basheer R. Sleep and Brain Energy Levels: ATP Changes during Sleep. Journal of Neuroscience. 2010;30:26:9007–16; doi:10.1523/jneurosci.1423-10.2010.

100. Kann O, Kovács R. Mitochondria and neuronal activity. American Journal of Physiology-Cell Physiology. 2007;292:2:C641–C57; doi:10.1152/ajpcell.00222.2006.

101. Rice D, Barone S. Critical periods of vulnerability for the developing nervous system: evidence from humans and animal models. Environmental Health Perspectives. 2000;108:suppl 3:511–33; doi:10.1289/ehp.00108s3511.

102. Bales KL, Saltzman W. Fathering in rodents: Neurobiological substrates and consequences for offspring. Hormones and Behavior. 2016;77:249–59; doi:10.1016/j.yhbeh.2015.05.021.

103. Curley JP, Champagne FA. Influence of maternal care on the developing brain: Mechanisms, temporal dynamics and sensitive periods. Frontiers in Neuroendocrinology. 2016;40:52–66; doi:10.1016/j.yfrne.2015.11.001.

104. Birrell JM, Brown VJ. Medial Frontal Cortex Mediates Perceptual Attentional Set Shifting in the Rat. The Journal of Neuroscience. 2000;20:11:4320–4; doi:10.1523/jneurosci.20-11-04320.2000.

105. Grissom NM, Reyes TM. Let’s call the whole thing off: evaluating gender and sex differences in executive function. Neuropsychopharmacology. 2018;44:1:86–96; doi:10.1038/s41386-018-0179-5.

106. Ribeiro S, Nicolelis MAL. Reverberation, storage, and postsynaptic propagation of memories during sleep. Learning &amp; Memory. 2004;11:6:686–96; doi:10.1101/lm.75604.

107. Boyce R, Glasgow SD, Williams S, Adamantidis A. Causal evidence for the role of REM sleep theta rhythm in contextual memory consolidation. Science. 2016;352:6287:812–6; doi:10.1126/science.aad5252.

108. Boyce R, Williams S, Adamantidis A. REM sleep and memory. Current Opinion in Neurobiology. 2017;44:167–77; doi:10.1016/j.conb.2017.05.001.

109. Abel T, Havekes R, Saletin Jared M, Walker Matthew P. Sleep, Plasticity and Memory from Molecules to Whole-Brain Networks. Current Biology. 2013;23:17:R774–R88; doi:10.1016/j.cub.2013.07.025.

110. Klinzing JG, Niethard N, Born J. Mechanisms of systems memory consolidation during sleep. Nature Neuroscience. 2019;22:10:1598–610; doi:10.1038/s41593-019-0467-3.

111. Jones MW, Wilson MA. Theta Rhythms Coordinate Hippocampal–Prefrontal Interactions in a Spatial Memory Task. PLoS Biology. 2005;3:12:e402; doi:10.1371/journal.pbio.0030402.

112. O’Neill PK, Gordon JA, Sigurdsson T. Theta Oscillations in the Medial Prefrontal Cortex Are Modulated by Spatial Working Memory and Synchronize with the Hippocampus through Its Ventral Subregion. Journal of Neuroscience. 2013;33:35:14211–24; doi:10.1523/jneurosci.2378-13.2013.

113. Alemany-González M, Gener T, Nebot P, Vilademunt M, Dierssen M, Puig MV. Prefrontal– hippocampal functional connectivity encodes recognition memory and is impaired in intellectual disability. Proceedings of the National Academy of Sciences. 2020;117:21:11788–98; doi:10.1073/pnas.1921314117.

114. Paul KN, Laposky AD, Turek FW. Reproductive hormone replacement alters sleep in mice. Neuroscience letters. 2009;463:3:239–43; doi:10.1016/j.neulet.2009.07.081.

115. Paul KN, Losee-Olson S, Pinckney L, Turek FW. The ability of stress to alter sleep in mice is sensitive to reproductive hormones. Brain research. 2009;1305:74–85; doi:10.1016/j.brainres.2009.09.055.

116. Ralston M, Ehlen JC, Paul K. Reproductive hormones and sex chromosomes drive sex differences in the sleep-wake cycle. Front Neurosci. 2024;18:1478820; doi:10.3389/fnins.2024.1478820.

117. Schwartz MD, Mong JA. Estradiol modulates recovery of REM sleep in a time-of-day-dependent manner. Am J Physiol Regul Integr Comp Physiol. 2013;305:3:R271–80; doi:10.1152/ajpregu.00474.2012.

118. Carone BR, Fauquier L, Habib N, Shea JM, Hart CE, Li R, et al. Paternally Induced Transgenerational Environmental Reprogramming of Metabolic Gene Expression in Mammals. Cell. 2010;143:7:1084–96; doi:10.1016/j.cell.2010.12.008.

119. Weaver ICG, Cervoni N, Champagne FA, D’Alessio AC, Sharma S, Seckl JR, et al. Epigenetic programming by maternal behavior. Nature Neuroscience. 2004;7:8:847–54; doi:10.1038/nn1276.

